# Evolution at two time-frames: ancient and common origin of two structural variants involved in local adaptation of the European plaice (*Pleuronectes platessa*)

**DOI:** 10.1101/662577

**Authors:** Alan Le Moan, Dorte Bekkevold, Jakob Hemmer-Hansen

## Abstract

Changing environmental conditions can lead to population diversification through differential selection on standing genetic variation. Structural variant (SV) polymorphisms provide examples of ancient alleles that in time become associated with novel environmental gradients. The European plaice (*Pleuronectes platessa*) is a marine flatfish showing large allele frequency differences at two putative SVs associated with environmental variation. In this study, we explored the contribution of these SVs to population structure across the North East Atlantic. We compared genome wide population structure using sets of RAD sequencing SNPs with the spatial structure of the SVs. We found that in contrast to the rest of the genome, the SVs were only weakly associated with an isolation-by-distance pattern. Indeed, both SVs showed important allele frequency differences associated with two different environmental gradients, with the same allele increasing both along the salinity gradient of the Baltic Sea, and the latitudinal gradient along the Norwegian coast. Nevertheless, both SVs were found to be polymorphic across most sampling sites, even in the Icelandic population inferred to originate from a different glacial refuge than the remaining populations from the European continental shelf. Phylogenetic analyses suggested that the SV alleles are much older than the age of the Baltic Sea itself. These results suggest that the SVs are older than the age of the environmental gradients with which they currently co-vary. Interestingly, both SVs shared similar phylogenetic and genetic diversity, suggesting that they have a common origin. Altogether, our results suggest that the plaice SVs were shaped by evolutionary processes occurring at two time-frames, firstly following their common origin and secondly related to their current association with more recent environmental gradients such as those found in the North Sea − Baltic Sea transition zone.

## Introduction

Disentangling neutral from selective forces acting on the process of population divergence is a key part of population genetics as the two forces provide different information about the mechanisms shaping population structure (Wright, 1978). While neutral markers provide data about population connectivity and demographic processes (Beaumont and Nichols, 1996), markers under selection can reveal the existence of locally adapted populations or cryptic speciation processes ongoing in a meta-population (Gagnaire *et al.*, 2015, Benestan *et al.*, 2018). Access to high numbers of genetic markers obtained from high throughput sequencing technologies has increased the power to detect genetic markers under selection. Recently, such genomic approaches have highlighted the importance of structural variants (SVs), which have often been associated with signals of selection in natural populations (see special issues: Wellenreuther *et al.*, 2019). SVs can originate from changes in copy number (deletion, insertion and duplication), orientation (inversion) or position (translocation, fusion) of DNA sequences resulting in reduced levels of recombination between rearranged parts of the genome (Kirkpatrick, 2010). Reduced recombination causes such chromosomal rearrangements to follow independent evolutionary pathways, often showing higher levels of population divergence than collinear regions of the genome (Navarro and Barton, 2003a, 2003b; Farré *et al.*, 2013). The SVs can harbour hundreds of genes and thus may have functional consequences for the organism (reviewed in Wellenreuther and Bernatchez, 2018).

Structural variants have been found to be important for promoting the evolution and maintenance of locally adapted populations in the face of gene flow. In a system of interconnected populations, gene flow results in the rapid homogenisation of genetic variation and is the main evolutionary force acting against the process of divergence (Slatkin, 1987). In the most extreme condition, when gene flow is too high, mutations that are favourable in a local environment are likely to recombine with maladapted genomic backgrounds, and can be lost through swamping effects (Lenormand, 2002). However, if two (or more) locally advantageous mutations are located within the same SV, the absence of recombination with adjacent, maladapted genomic backgrounds increase their fitness locally. In comparison to independent genetic variants, mutations within SVs are less affected by the migration load and therefore more likely to maintain locally adapted alleles in a meta-population (Kirkpatrick and Barton, 2006). The strength of population differentiation will depend on the balance between genetic drift, migration and selection, the effect of the latter being inflated if both locally adapted variation and genetic incompatibilities (decreased fitness in heterozygotes) are found at the same time in the population or even within the same SV (Bierne *et al.*, 2011; Faria *et al.*, 2019). Co-adaptation involving positive epistatic interactions between loci is also more likely to be maintained within a SV (Dobzhansky, 1970; Feldman *et al.*, 1996), leading to the development of a supergene that further increases the adaptive potential of SVs (Thompson and Jiggins, 2014).

Structural variants have often been found to be an order of magnitude older than the age of the populations in which they are found, suggesting that their adaptive potential often relies on ancient polymorphisms (reviewed in Wellenreuther and Bernatchez, 2018; Marques et al., 2019), in effect representing a source of standing variation for population divergence and adaptation. For instance, million years old SVs have promoted the repeated evolution of ecotypes following the post glacial recolonization of new environments, as described in systems undergoing parallel evolution (e.g. Jones *et al.*, 2012; Nelson and Cresko, 2018; Morales et al., 2019). It has been suggested that such adaptive genetic variation can be shaped during distinct time periods (i.e. “Evolution at two time-frames”, c.f. Van Belleghem *et al.*, 2018). The first period corresponding to the initial increase in frequency of adaptive variations in the absence or under limited gene flow, while the second period is associated with their sometimes much later association with environmental, physical and/or endogenous barriers to gene flow, observable in contemporary meta-populations (Bierne *et al.*, 2011, Van Belleghem *et al.*, 2018). While the origin of a SV often remains unclear, they sometimes originate from adaptive introgression of loci that have positive fitness effects for the introgressed species or population. For example, adaptive introgression of a major inversion is responsible for mimicry of the wing colour pattern between poisonous *Heliconius* butterflies (The Heliconius Genome Consortium *et al.*, 2012; Jay *et al.*, 2018).

The European plaice (*Pleuronectes platessa*) is a marine flatfish in the eastern Atlantic found from the Iberian Peninsula in the South to the Barents Sea and Greenland in the North. This species is thought to have colonized the northern range of its distribution (from the North Sea to Greenland) after the last glacial maximum (Hoarau *et al.*, 2002), and has also large populations established in the western parts of the Baltic Sea, a brackish environment formed eight thousand years ago (8 kya), which represents a low salinity environment for marine species (Johannesson and André, 2006). The biological traits of plaice are generally expected to be associated with low levels of population divergence (Ward *et al.*, 1994; Waples and Gaggiotti, 2006;): large effective population size (reducing genetic drift), external fertilization, non-determinant spawning season, pelagic egg and larval phases (promoting gene flow) (Harding *et al.*, 1978; Rijnsdorp, 1991). Previous studies based on six microsatellite loci have found weak, often statistically non-significant, levels of population structure across Europe, except across the bathymetric barrier between the continental shelf and off-shelf regions (Europe vs Iceland and Faroe Islands, respectively), where depth apparently acts as a strong physical barrier (Hoarau *et al.*, 2002; Was *et al.*, 2010). However, the off-shelf population shows a higher genetic diversity than most of the populations from the northern distributional range of the plaice (Hoarau *et al.*, 2002), which could suggest that this population has a more ancient origin than what is currently thought. In a recent study of the genomic basis underlying the colonization of the Baltic Sea, we identified two large polymorphic SVs (Le Moan *et al.*, 2019) carrying a strong signal of selection between the North Sea and the Baltic Sea plaice populations (66% of the top 0.1% outlier loci localized within two SVs). However, whether these SVs evolved in connection with a founder event and adaptation to a Baltic Sea environment or are ancient polymorphisms remained unresolved. The European plaice is known to hybridize with its sister-species, the European flounder *(Platichthys flesus)* (Kijewska *et al.*, 2009). The flounder is a euryhaline species better adapted to low salinity than the plaice, and can be found in freshwater lakes and in the innermost parts of the Baltic Sea where plaice does not occur (Hemmer-Hansen *et al.*, 2007). Therefore, it is possible that the European flounder is the source of the SVs found in plaice.

In the present study, our overall objective was to increase our understanding of the origin of the SVs and their effects in contemporary populations of the European plaice. Specifically, the main goals of the study were to: i) test for a potential flounder origin of SVs; ii) estimate the age of the SVs in plaice with a phylogenetic approach; iii) re-assess the population structure in European plaice from northern Europe and from Iceland with the use of a population genomics approach, iv) evaluate the contribution of SVs to population structure, and v) provide relevant data to understand the extent to which selection is involved in maintaining the allelic clines observed at the SV. This work thus provides increased insight into the relative roles of environmental gradients, demographic history, hybridization and genomic structural changes in the evolution of a widespread and highly abundant species.

## Materials and methods

### Geographical sampling

European plaice samples were collected at seven sites distributed across Northern Europe and Iceland (Figure 1a & Table 1) during the spawning season (Rijnsdorp, 1991). Samples from the North Sea, Kattegat, the Belt Sea and the Baltic Sea were collected in 2016-2017. These samples were also the subject of the study by Le Moan *et al.* (2019) that studied the diversification process involved in the colonization of the Baltic Sea across multiple flatfish species. In that study, two large SVs were identified with allele frequency differences associated with the salinity gradient between the North Sea and the Baltic Sea. To explore the spatial distribution of these SVs in greater detail, three additional northern sites were included in the current study. These samples were collected from the Barents Sea, Norway and Iceland in 2013. Most of the northern parts of the plaice distribution was covered with this sampling design. Analyses included 255 plaice in total, along with ten European flounder (the species hybridizing with plaice in the study area), and ten common dab (*Limanda limanda*), a closely related but reproductively isolated species from both European plaice and European flounder, to be used as outgroup in the phylogenetic analyses.

**Table 1:** Details of the sampling locations, with the corresponding site ID, sample sizes, longitude, latitude and the date when the samples were collected.

| Location | Site ID | Sample size | Sample analysed | Longitude | Latitude | Dates of sampling |
| --- | --- | --- | --- | --- | --- | --- |
| Iceland | Iceland | 25 | 25 | 62.9 | -15.8 | Feb. 2013 |
| Barents Sea | Barents Sea | 25 | 19 | 70.7 | 20.2 | Feb. 2013 |
| Norway | Norway | 25 | 24 | 64.8 | 8.8 | Feb. 2013 |
| North Sea | North Sea | 30 | 26 | 55.26 | 7.09 | Feb. 2016 |
| Kattegat - Skagerrak | Kattegat | 50 | 44 | 57.63 | 9.30 | Feb. 2017 |
| Øresund-Belt Sea | Belt Sea | 50 | 48 | 54.65 | 10.64 | Mar. 2016-2017 |
| Southwest Baltic | Baltic Sea | 50 | 48 | 54.98 | 14.54 | Mar. 2017 |

**Figure 1:**
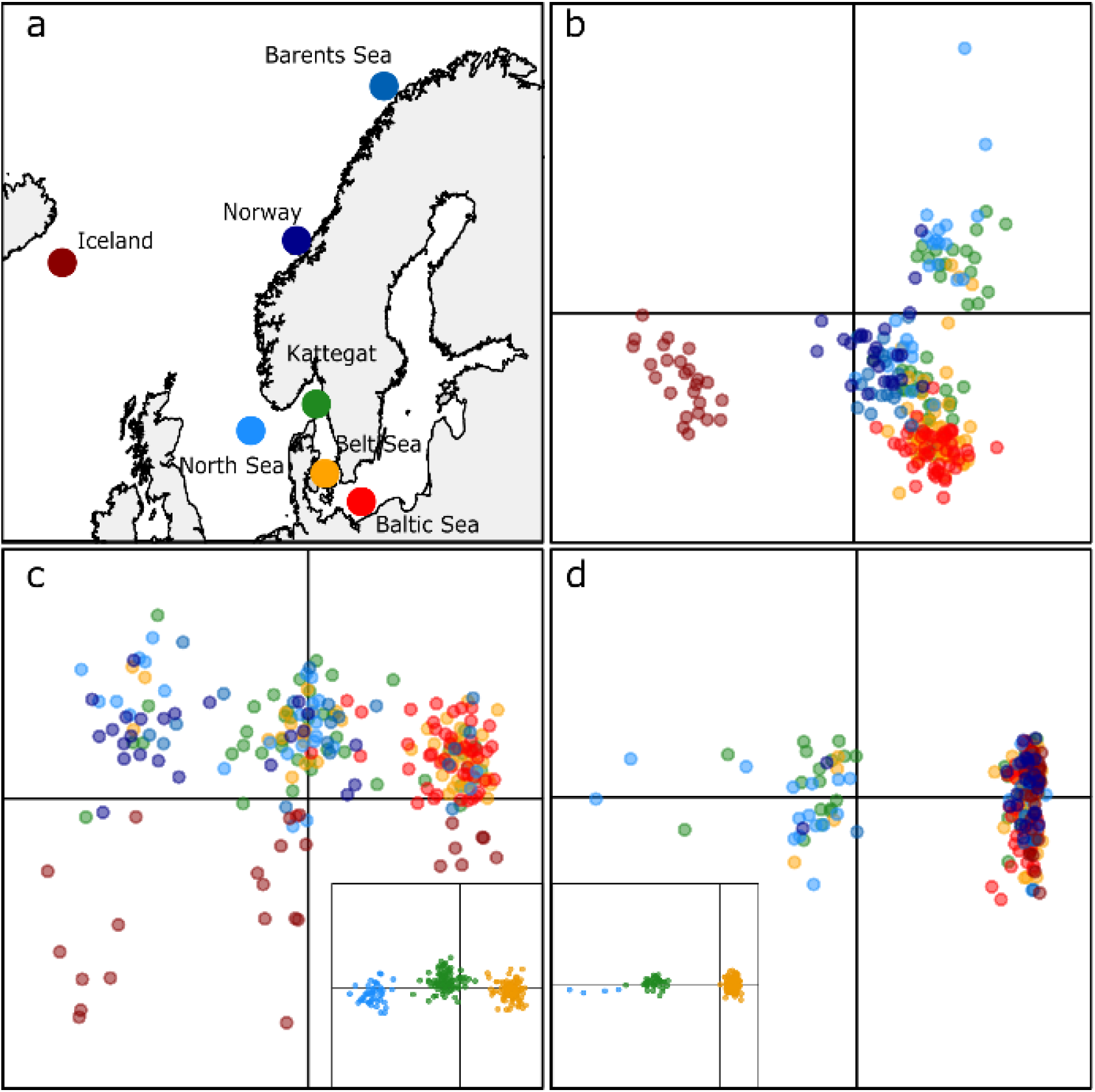
Sampling design (a) and principal component analyses performed on individual diversity for the European plaice based on a genomic dataset of 3019 SNPs in the overall dataset (b), 222 SNPs on chromosome 19 (c) and 210 SNPs on chromosome 21 (d). Colours correspond to sampling sites represented on the map in a). Inserts in c) and d) show DAPC results grouping individuals by haplogroup, where green dots are individuals from the heterozygote haplogroup and the yellow and blue dots are individuals from the homozygote haplogroups 1 and 3, respectively.

### ddRAD libraries and sequencing

Whole genomic DNA was extracted from gill tissue using the DNeasy Blood Tissue kit (Qiagen). The DNA concentration was measured with the Broad Range protocol of Qubit version 2.0® following the instructions manual. DNA extractions were diluted to 20 ng/µl. Four double-digest RAD (ddRAD) libraries were constructed following Poland and Rife (2012), using Pst1 and Msp1 restriction enzymes with rare and frequent cutting sites, respectively. Each library was made by randomly pooling between 60 and 75 barcoded individuals from various locations. The libraries were size-selected on agarose gels in order to retain insert sizes between 350 and 450bp. After an amplification step (12 cycles), the libraries were purified with AMPure® beads and their quality was checked on a Bioanalyzer 2100 using the High Sensitivity DNA protocol (Agilent Technologies). The targeted size selection was successful in most libraries except in one that was slightly shifted towards shorter insert size (from 300 to 400 bp). The effects of different library size ranges are discussed further below. Each library was pair-end sequenced on one Illumina HiSeq4000® lane (2*101 bp).

### Bioinformatics

Raw sequences were processed using the “ref-map” pipeline from Stacks version 2.1 (Catchen *et al.*, 2013). Specifically, the samples were demultiplexed with “process radtag” by removing reads with mean sequencing quality below 10 and reads with uncalled base pairs. On average, we obtained six million reads per sample (Figure S1). The reads were trimmed to 85 bp using trimmomatic (Bolger *et al.*, 2014), and aligned to the Japanese flounder (*Paralichthys olivaceus*) genome (Shao *et al.*, 2017) using bwa-mem set with default parameters (Li and Durbin, 2009). This reference genome is from a species of the same family of the European plaice, which has a relatively conserved genome structure (Robledo *et al.*, 2017). On average, 65% of reads per sample mapped to the reference genome (Figure S1). SNPs were called based on the mapping results using the “gstacks” function with the “marukilow” model set with a minimum of coverage (m) of 5X to build a stack of identical reads into a biological sequence, and alpha parameter of 0.05. Only bi-allelic SNPs genotyped in at least 80% of the individuals within each sampling site and with a maximum heterozygosity of 0.80 were called using the “population” function. All individuals with more than 10% missing data were removed (details in Table 1). Finally, SNPs with a significant departure from Hardy-Weinberg equilibrium (p-value 0.05) in more than 60% of the sampling sites, as well as singletons, were removed using vcftools (Danecek *et al.*, 2011). The average coverage after filtering was 29X per sample (Figure S1). Unfortunately, size selection was slightly shifted between the first three libraries (containing North Sea and Baltic Sea samples) and the last library (containing Barents Sea, Norway and Iceland samples). This shift resulted in a reduction of the genomic sampling when keeping only loci sequenced for all the sampling sites. Therefore, three datasets were constructed to take the differences in size selection into account: the “overall dataset” including all sampling sites (8587 RAD-tags with 28,016 SNPs genotyped in 234 individuals), the “southern dataset” including the North Sea, Kattegat, Belt Sea and Baltic Sea (17,342 RAD-tags with 56,740 SNPs genotyped in 166 individuals), and the “northern dataset” including Iceland, Norway and Barents (35,549 RAD-tags with 92,706 SNPs genotyped in 68 individuals). These data sets were subsequently used for different analyses focusing on different aspects of population differentiation (see below), and further filtered to fulfil the requirements of the analyses performed. More details about the number of RAD-tags and the number of SNPs obtained after each filtration step can be found in Table S1.

### Population structure and demographic history

The “overall dataset” was used to describe overall population structure and was thinned by removing loci with minor allele frequencies (MAF) below 5% and by keeping only one random SNP per bin of 1kb to limit effects from physical linkage but not the Linkage Disequilibrium (LD) within SVs (Figure S9). In total, we kept 3019 SNPs from which individual genetic diversity was visualized using PCA analyses, conducted with the R package adegenet (Jombart, 2008). The same package was used to compute population specific heterozygosity. Pairwise genetic differentiation (*F*_ST_) between samples was estimated following the method of Weir and Cockerham (1984) using the R package StAMPP (Pembleton *et al.*, 2013). We used 1000 bootstraps over loci to evaluate if pairwise *F*_ST_ values were significantly different from 0. The effect of isolation-by-distance was assessed with a Spearman correlation test between pairwise *F*_ST_ and geographical distances among sampling sites and assessed with a Mantel test set after 9999 permutations with the R base package (Ihaka and Gentleman, 1996). All analyses were conducted by including and excluding the two chromosomes carrying the SVs, as well as using the information from these two chromosomes only. Additionally, the correlation between genome-wide population structure and the structure displayed by the structural variants was assessed by Mantel tests on the genome-wide *F*_ST_ matrix without the SVs and both *F*_ST_ matrices of the structural variants only.

We used an approximation-of-diffusion approach, as implemented in the software δaδi (Gutenkunst *et al.*, 2010), to examine the demographic histories associated with the major population breaks that separate populations from the continental shelf and Iceland (Hoarau *et al.*, 2002). Specifically, we assessed the demographic history of the divergence of Iceland and its closest continental shelf population sample from Norway, using the “northern” dataset. For this analysis, only polymorphic sites with a minimum allele count of 2 in at least one of the two populations were kept, which were then filtered for LD (1 SNP per 1000kb). In total, four standard scenarios of demographic history were compared: the Strict Isolation (SI), the Isolation-with-Migration (IM), the Ancestral Migration (AM), and the Secondary Contact (SC) models (Fig. S9). These models were compared to the data using the folded version of the Joint Allelic Frequency Spectrum (JAFS) without considering singletons (-z option), using a modified version of δaδi from Tine *et al.* (2014). The best model was then selected based on its goodness of fit using the Akaike Information Criterion (AIC) selection.

### Genomic analyses of the structural variants

The two chromosomes carrying the putative SVs (named respectively, C19 and C21 and SV19 and SV21 in the following), were extracted from the overall dataset (LD and MAF pruned) to construct two independent sub-datasets to examine population structure in the SVs alone, using a PCA approach (on 222 and 210 SNPs, respectively). Initial PCA plots showed clustering into three distinct groups (Figure 1) and hence suggested that each SV behaves like a Mendelian character with two major divergent alleles of multiple linked loci leading to three distinct “haplogroups” (two homozygotes and one heterozygote). Consequently, we performed DAPC analyses with adegenet (Jombart and Ahmed, 2011) set to three groups, to identify the haplogroup of each individual using the find.cluster function. The allele frequencies of the SVs for each population were calculated based on the DAPC clusters, using the formula

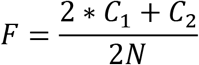

Where C_1_ is the number of individuals assigned to one of the homozygote haplogroups (haplogroup 1 or 3) and C2 the number of individuals assigned to the heterozygote haplogroup (haplogroup 2) in the DAPC, and N is the number of samples in the population. Several SNPs showed observed heterozygosity (H_O_) of 1 in haplogroup 2 (Figure S6) which also had *F*_ST_ = 1 between haplogroup 1 and 3 carrying different alleles (Figure 3c-d). These results are expected for SVs and therefore validated our genotyping procedure. Then, we used the DAPC groups as genotype input to calculate the pairwise *F*_ST_ between populations at each SV using hierfstat (Goudet, 2005).

We analysed the subsets of the overall data (“northern” and “southern” dataset combined) without pruning for LD to increase the genomic coverage and better describe the genomic heterogeneity along the chromosomes carrying SVs. Nucleotide diversity (π) and H_O_ were calculated per SNP for each haplogroup inferred in the DAPC. The genomic distribution of differentiation was examined using SNP specific *F*_ST_ values. Specifically, this differentiation was calculated for Norway vs. Barents Sea, Norway vs. Iceland using the northern dataset, and for North Sea vs. Baltic Sea and between the homozygote haplogroups 1 and 3 using the southern dataset. We used a quantile regression from the quantreg R package to calculate the variation of the upper 1% *F*_ST_ quantile and average *F*_ST_ along bins of 100kb along the chromosome in the different pairwise comparisons. We focused on the upper quantile to limit effects from variable levels of diversity across the SVs that would tend to depress average *F*_ST_ in certain regions along the SVs (Figure S7). Finally, we estimated LD by calculating the pairwise correlation between loci localised on chromosomes 19 and 21. These statistics were computed using vcftools (Danecek *et al.*, 2011) and the smoothed value of the statistics were performed with ggplot2 using local regression with loess (Wickham and Winston, 2008) with a span parameter of 0.3. In addition, we used LDheatmap (Shin et al., 2006) that integrates the information about the physical position of the SNP on the chromosome to represent the heatmap of LD between every pair of SNPs.

### Gene content of the structural variants

In order to understand the genetic composition of the two SVs, we extracted the filtered RAD-tags from the two SVs into two individual fasta files. The genomic ranges of the structural variants were defined based on visual observation of the breaking point positions and on the *F*_ST_ values between homozygote haplogroup 1 and 3 individuals, starting after the first and ending before the last SNP with *F*_ST_ > 0.9 (from 1.4 to 9.9 Mbp for SV19 and from 10.5 to 20 Mbp for SV21). The RAD-tags within the SVs were then aligned back to a previous version of the Japanese flounder genome that is annotated (Shao *et al.*, 2017) using bwa-mem set with default parameters (Li and Durbin, 2009). All the genes localized between the two most distantly aligned RAD-tags on the genome were then extracted from the annotation file using custom bash scripts. Finally, the gene lists were mapped to the Gene Ontology resource website (http://geneontology.org/) to check for functional enrichment using the zebra fish and the human databases. The functional enrichment analyses were performed using both gene lists independently and with the gene lists combined.

### Phylogenetic analyses

The ddRAD protocol can be used to identify orthologous sequences with restriction sites conserved across distantly related species. As such, this property was used in order to estimate the age of the SV polymorphisms by building a phylogeny of European plaice and two other species of the Pleuronectidae, the European flounder and the common dab. In order to obtain the sequences of the homozygote haplogroups and infer their divergence, we only retained the haplogroup 1 and 3 plaice individuals (based on the DAPC) from the southern dataset. We focused on the southern dataset because it was the only dataset including haplogroup 3 for SV21 (Figure 1d) and with the highest number of reads overlapping between the plaice and the two outgroup species.

Three independent phylogenies were constructed using concatenated ddRAD loci, one representing each SV and one representing loci localized outside the SVs. The loci from the SVs were extracted based on the genomic ranges defined above for gene content analyses, while the loci representative of the genome-wide divergence were selected from outside the SVs to get a sequence length similar to that of the SVs (from 15 and 25 Mbp on chromosome 19). Chromosome 19 was used to infer phylogenies both within the SV and to represent a genome wide phylogeny in order to estimate any effects that the SVs may have on collinear regions of the same chromosome. Only ddRAD loci with sequence information for both homozygote haplogroups in plaice and in flounder and/or dab were used for the phylogeny. The full sequence of each RAD locus was extracted into individual fasta files with the population function of Stacks (Catchen *et al.*, 2013) using a “whitelist” (-w) option comprising the RAD-tag ID from the filtered southern dataset. For each RAD locus, one random RAD allele per individual was retained. All alleles were concatenated into one pseudo-sequence using a custom script. The three phylogenies were inferred based on orthologous sequences of 10125 bp, 6152 bp and 10341 bp for chromosomes 19, 21 and genome-wide, respectively. All phylogenies were estimated in RaxML (Stamatakis, 2014), under the GTR+GAMMA model with a random number set as seed. Finally, we tested for potential gene flow between species using the “f4” statistic from Treemix (Pickrell and Pritchard, 2012), evaluating the mismatch between the tree topology inferred with RaxML and individual SNP topologies.

The length of the inferred branch between each cluster represents the number of substitutions occurring after the split of the species/haplogroups, and it is directly proportional to the time of divergence under neutral processes (Kimura, 1983). We applied a strict molecular clock to transform this nucleotide divergence into time since divergence in years. Specifically, we divided the substitution rate along each branch by the average SNP mutation rate (10^−8^; Tine *et al.*, 2014), and multiplied this value by 3.5, the average generation time of the Pleuronectidae (Erlandsson *et al.*, 2017). Although this approach is commonly used to age SVs, the divergence estimates should be interpreted with caution as cross recombination events in the centre of the SV would decrease the estimate of divergence and selection acting on the haplogroup (background and disruptive) would tend to increase the estimate of divergence.

## Results

### Population structure and demographic history

The first axis of the PCA explained 1.45% of the total inertia and distinguished Icelandic plaice from all continental shelf individuals, and to a lower extent also identified separation of (marine) Atlantic samples from (brackish) Baltic Sea samples (Figure 1b). The second axis explained 1.2% of the inertia that roughly traced the gradient from the North Sea to the Baltic Sea. Observed heterozygosity was maximal in the North Sea (H_O_ = 0.193/0.190 with/without SVs) and decreased across the North Sea-Baltic Sea transition zone (Kattegat H_O_ = 0.185/0.184, Belt Sea H_O_ = 0.182/0.184 and Baltic Sea H_O_ = 0.178/0.181) as well as towards the north (Norway H_O_ = 0.180/0.180 and Barents Sea H_O_ = 0.179/0.180). The H_O_ of the Icelandic population was intermediate (H_O_ = 0.184/0.185). All pairwise *F*_ST_ estimates were significantly different from zero, except for the two sites in the North Sea-Baltic Sea transition zone (Kattegat vs Belt Sea, Table S2). Pairwise comparisons including the Iceland population were homogeneously valued around *F*_ST_ = 0.03, (Figure 2, yellow dots) while all other pairwise *F*_ST_ estimates were lower (Figure 2, red dots). The demographic modelling revealed that the most likely scenario for the origin of the separation between Iceland and the continental shelf was a scenario of past isolation followed by a secondary phase of gene flow (SC model – Figure S11 and Table S5). The difference in AIC between this model and the second best model (Ancestral migration; AM) was 2912, which represents a very strong support for the SC among the tested models (Rougeux *et al.*, 2018). The estimate of the time of divergence, including the isolation phase, was ten times higher than estimated time under the secondary contact phase (Table S5). By using a generation time of 3.5 years and a mutation rate of 10^−8^ per generation, the time of split was estimated to approximately 112 kya (secondary contact estimated to 9 kya).

**Figure 2:**
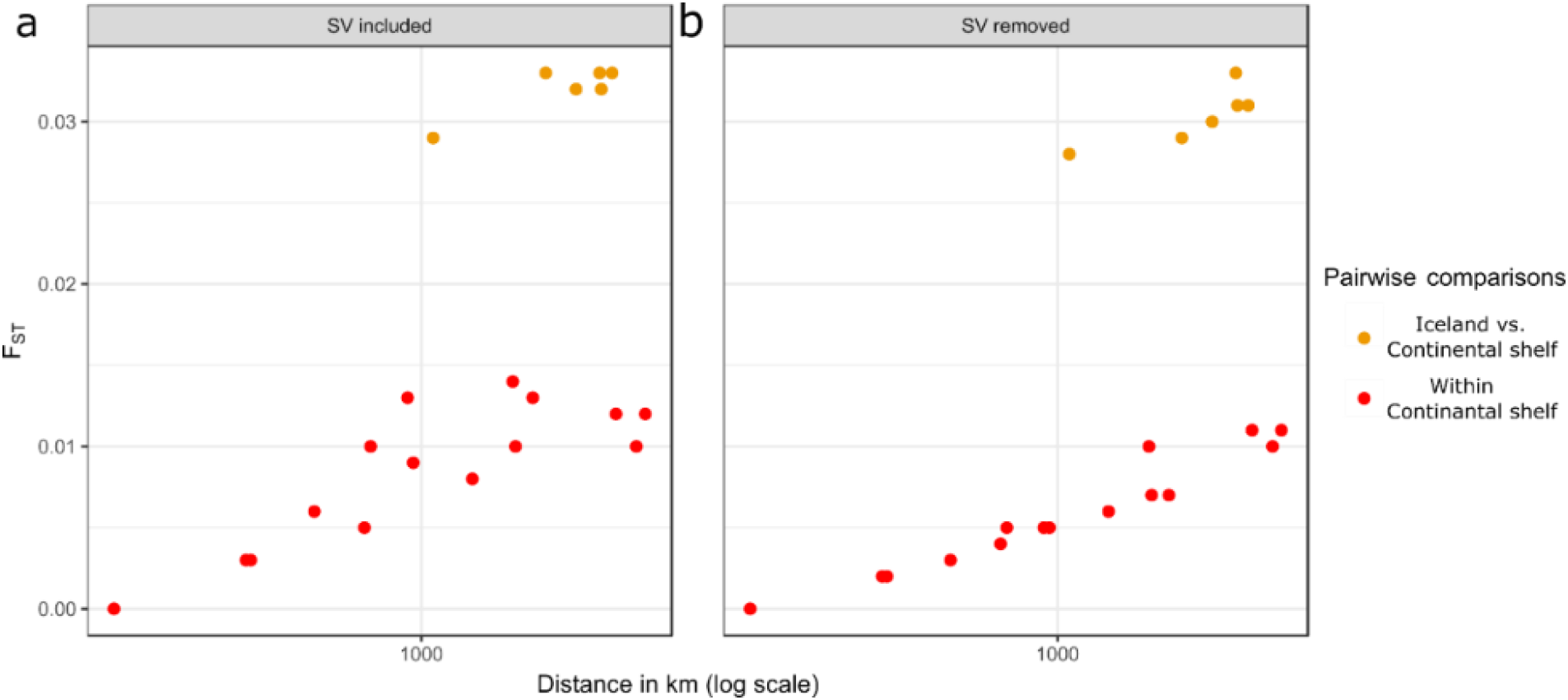
Relationships between geographic distance and genetic differentiation, including structural variants in a (r=0.56) and excluding structural variants in b (r=0.96). Comparisons including the Icelandic sample (yellow dots) were excluded from the correlation analyses.

**Figure 3:**
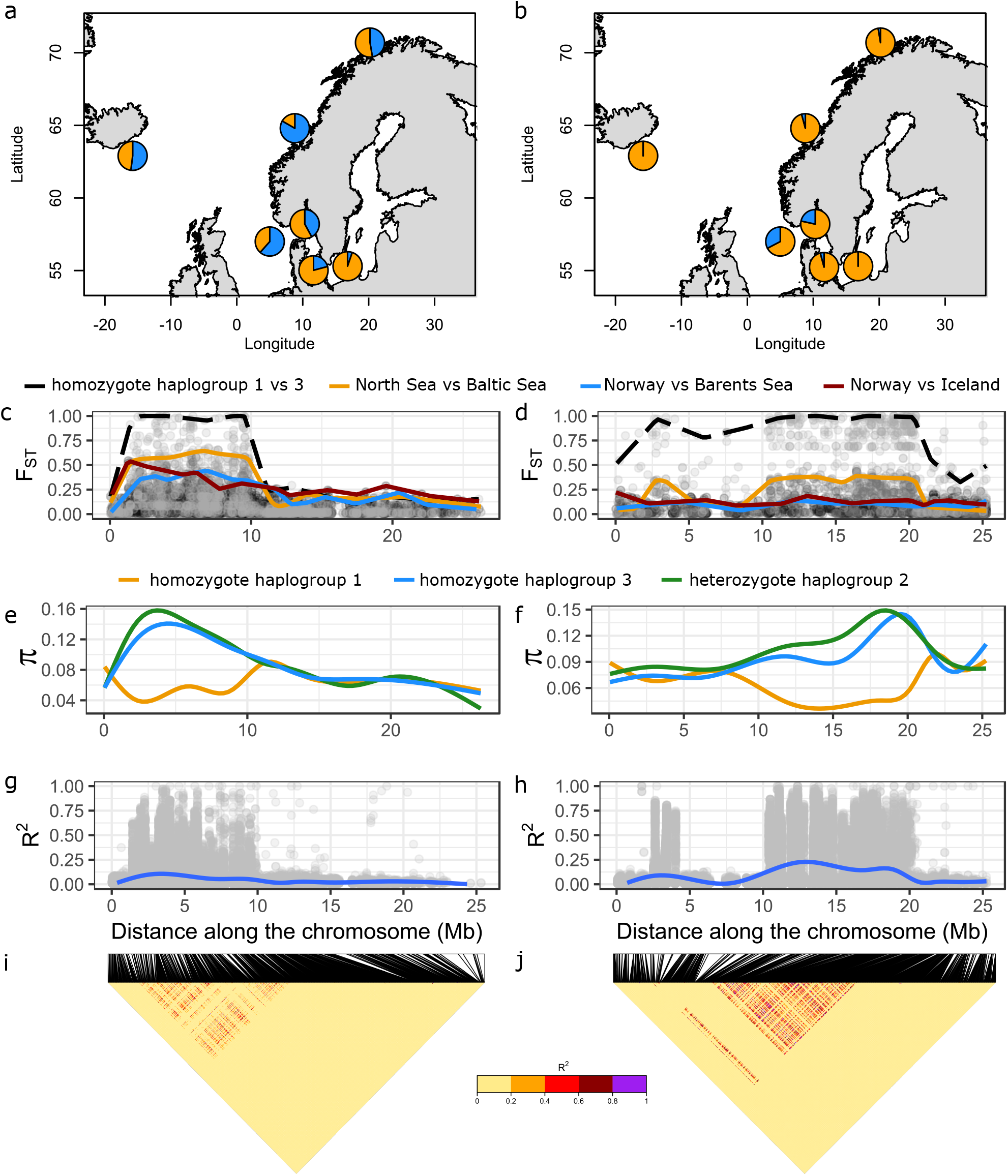
Sample specific data for the European plaice structural variants SV19 (panels a, c, e, g, i) and SV21 (panels b, d, f, h, j): a-b. sample specific allele frequencies; c-d. differentiation between different pairs of populations and between haplogroups 1 and 3 along the two chromosomes carrying the SVs, each dot is the FST value of an individual locus, and the lines represent the smoothed upper 1% quantile (one colour = one comparison, alpha =0.3); e-f. smoothed average for the three haplogroups identified in the DAPC (span = 0.3); g-h. variation of LD along the chromosomes and i-j. LD heatmaps for pairwise SNP comparisons.

In order to exclude effects from ancient demography on inference about contemporary neutral structure, the Icelandic population was removed from the analyses of isolation-by-distance. The pairwise *F*_ST_ across the continental shelf was significantly correlated with geographical distance (Mantel test: r = 0.59, p > 0.01). Interestingly, removing the structural variants from the analysis reduced the variation of the pairwise *F*_ST_ (Figure 2, red dots) and resulted in a stronger correlation between genetic and geographic distances (r = 0.96, p > 0.01). However, this pattern of isolation-by-distance was not detected when only the chromosomes carrying SVs were analysed (r = 0.06, p = 0.89 for C19 & r = 0.11, p = 0.35 for C21, Figure S2 & Figure S3). Consequently, the genome-wide pairwise *F*_ST_ (without the structural variants) was not correlated with the pairwise *F*_ST_ calculated with the chromosomes carrying SVs (r=0.08, p=0.77 for C19, & r=0.11, p=0.64 for C21). Altogether, this result suggests that the population structure found genome wide is different from the population structure carried by the SVs. Interestingly, the pairwise *F*_ST_ values calculated for the two chromosomes with structural variants were also not significantly correlated (r=−0.2, p=0.38), which suggests unique geographical patterns of structuring for the two variants.

### Genomic variability and differentiation of the structural variants

A high dispersion of samples from the North Sea and Kattegat was observed on the second axis of the PCA (light blue and green samples in Figure 1b). This dispersion mostly involved SNPs from chromosome 19 and 21 carrying structural variants (Figure S4b showing the PCA loading plot). The PCA based on these chromosomes showed three clusters of samples which were also inferred in the DAPC analyses (Figure 1c & d). In both cases, the clusters were likely due to the presence of two divergent multi-locus alleles segregating in the plaice populations for each SV, resulting in three haplogroups corresponding to two homozygotes (haplogroups 1 and 3 in blue and yellow, respectively) and one heterozygote cluster (haplogroups 2, in green).

Both SVs were polymorphic at most sampling sites (Figure 3a-b). However, whereas SV19 allele frequencies were variable across most sampling sites (Figure 3a), only the North Sea and Kattegat showed both SV21 alleles in high frequency (Figure 3b and Figure S5), which confirmed the different geographical patterns for the two SVs also identified by a lack of correlation of pairwise *F*_ST_ estimates. The important variation in allele frequencies leads to high *F*_ST_ between population that are geographically close (Table S4, illustrated Figure 3c). This large differentiation was evident across nearly one half of the chromosomes, from 1.4 Mbp to 9.9 Mbp for SV19 (Figure 3c), and from 10.5 to 20 Mbp for SV21 (Figure 3b). The genome-wide differentiation outside the SVs was lower (e.g. mean *F*_ST_ North Sea vs Baltic Sea= 0.004, sd=0.031) than inside the SVs (mean *F*_ST_ SV19= 0.203, sd=0.177 & mean *F*_ST_ SV21=0.167, sd=0.142; Figure 3b and Figure S10). The individuals from haplogroups 1 and 3 were differentially fixed for eight and 30 SNPs within SV19 and SV21, respectively (also represented by the black dashed line in Figure 3b). Strong LD occurred along the entire SVs, confirming that recombination between the SV alleles is rare (Figure 3d). Only haplogroup 1 in each SV (in yellow in Figure 3) showed reduced genetic diversity whereas haplogroup 3 (in blue in Figure 3) had a higher genetic diversity than the average observed across the genome. As expected, the individuals from haplogroup 2 showed the highest diversity of the three DAPC groups. The two SVs showed higher levels of linkage disequilibrium than the average genome-wide (0.2 vs. 0.07), but lower levels than the average LD within the SVs (Figure S8). Chromosome 21 showed two peaks of LD separated by a distance of 5Mbp of limited LD (Figure 3d). The LD within the two regions were equivalent to the LD between these regions (Figure 3d).

### Gene content and functional enrichment within the structural variants

We identified 200 and 256 genes located within SV19 and SV21, respectively (Supplementary file I). Of these genes, 48/66 in SV19/SV21 had a known function in zebra fish, and 145/191 in humans. For SV21, we detected functional enrichment linked to physiological or morphological membrane disruption (>100% enrichment, p = 3.1×10^−10^), immune and defence response to bacteria (16%, p = 1,6×10^−05^), and chromosome/chromatin organization (10%, p = 6.5×10^−6^) using the zebra fish database, and for purinergic receptor (40%, p=4.52×10^−7^) using the human database. However, no functional enrichment was identified for genes in SV19. By pooling both gene lists only the gene enrichment for SV21 was detected without additional function using the zebrafish database. However, using the human database, we identified functional enrichment for pre-B cell allelic exclusion (62%, p=7.5×10^−5^), which are genes involved in the immune system. Moreover, when looking specifically for known candidate genes involved in the local adaptation in other marine fishes, we found the presence of two genes linked to heat shock proteins (heat shock factor-binding protein 1 and 4) within SV19.

### Phylogenetic analyses

The common dab was the most divergent species in all phylogenetic trees, with 0.0240-0.0250 substitutions per site compared with European flounder and European plaice (Figure 4), which corresponds to time of split at 9 million years ago (Mya). All plaice individuals were equally distant from flounder individuals based on the concatenated ddRAD loci representative of the genome wide divergence, with an average of 0.0110 substitutions per site (4 Mya, Figure 4). However, the two haplogroups were clearly divergent in both SV phylogenies, with an average distance of 0.0020 and 0.0017 substitutions per site for SV19 and SV21 (Figure 4), respectively. In each case, the deepest branches were observed for haplogroup 1 (yellow) which lead to a different estimate of divergence for each branch, around 460 (504-434) and 200 (224-179) kya for haplogroups 1 and 3, respectively. The longest branch of the SVs resulted in an average plaice-flounder divergence slightly higher than outside the SVs (0.0120 and 0.0115 substitutions per site, respectively).

**Figure 4:**
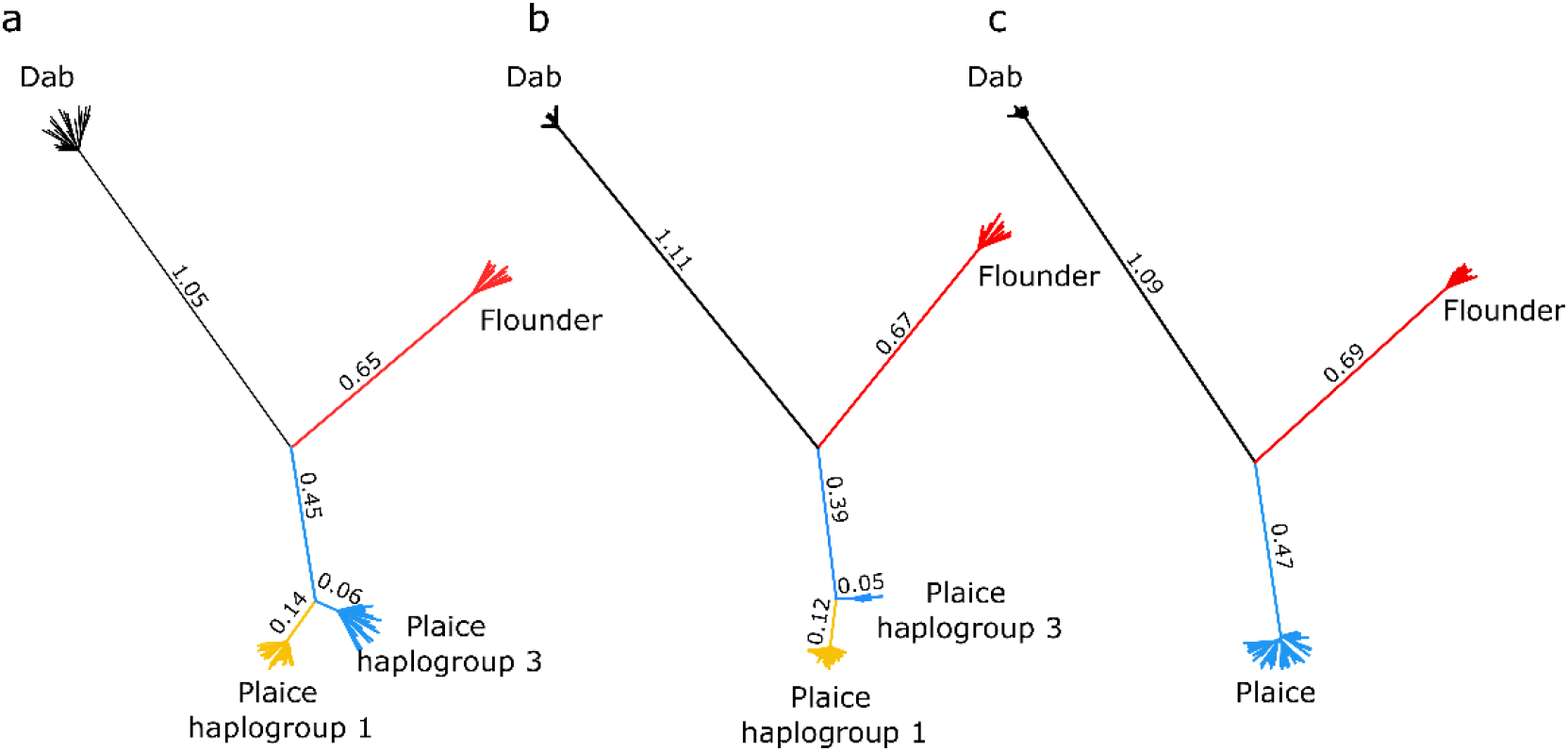
Phylogenetic trees for the ddRAD loci within SV19 (a), SV21 (b) and outside the SVs (c). The lengths of the branches reflect the numbers of substitutions per site (multiplied by 100 in the figures).

We found a low, but statistically significant, effect of introgression between the plaice and the flounder (f4 = 0.004, p-value < 0.05). However, this signal was mostly carried by three loci from SV19 and one from SV21 that showed a departure with the tree phylogeny of the structural variants. In all cases, the flounder and haplogroup 1 of the plaice were nearly fixed for the same allele, which was different from the major allele observed in the common dab and within haplogroup 3 of the plaice. Removing these SNPs lead to f4 statistic values not significantly different from 0 (i.e. indicating no effect of introgression).

## Discussion

The two SVs previously identified in Le Moan *et al.* (2019) along the North Sea − Baltic Sea transition zone were polymorphic across most of the north-eastern Atlantic distribution range of European plaice (Figure 3 a-b). Our analyses confirm the clear isolation of the Icelandic population and add new perspectives to our understanding of the evolutionary history of European plaice by identifying this population as likely originating from a different glacial refugium than the continental shelf populations. This knowledge has important implications for our understanding of the origin and evolution of structural variants in the species. We have also confirmed weak but significant genetic structure among continental shelf samples. Removing the SVs from the analyses leads to a stronger pattern of isolation-by-distance among European plaice samples (from r = 0.56 to r = 0.96) which was not detected in previous European plaice studies based on six microsatellite markers (Hoarau *et al.*, 2002; Was *et al.*, 2010). Thus, the results highlight the power of increasing genomic resolution to detect subtle population structure for species with high gene flow. Below, we discuss the evolution and effects of structural variants in European plaice in the context of population history and structure of the species in the northern Atlantic.

### Origin of structural variants in European plaice

The low genetic diversity of haplogroup 1 (yellow, Figure 3c) and their long branches in the phylogenies (Figure 4) suggest that they are the derived form of the SVs. The low diversity along haplogroup 1, and its association with the edge of the plaice distribution, suggest that these SVs are under some type of selective pressure, which is further supported by the deep branch inferred in the phylogenies.

#### Type of the structural variants

The two SVs covered nearly half of chromosomes 19 and 21 of the Japanese flounder genome, where a strong LD was maintained over 9Mbp. These large linkage blocks are expected with chromosomal rearrangements such as inversions, duplications and translocations which can formally be distinguished by use of a linkage map or genome sequencing (e.g. Faria *et al.*, 2019), but not with the reduced representation approach used in this study. However, our data filtration steps (filtering by heterozygosity) would have resulted in the loss of duplicated regions. Moreover, assuming a high degree of synteny between European plaice and Japanese flounder genomes, the size of the LD blocks and their central position are consistent with the presence of at least two major inversions in the genome (Kirkpatrick *et al.*, 2010). The observed second peak of LD on chromosome 21 could be due to the presence of a third small inversion. However, the similar value of LD within and between the two LD blocks on this chromosome suggest that they may be part of the same inversion and that a lack of synteny between the plaice and the Japanese flounder reference genome results in two distant peaks of LD on chromosome 21.

#### Age of the structural variants

The inferred derived allele of both SVs were found in both Iceland and continental shelf samples, presumably representing different glacial refugia, which were estimated to have diverged 112 kya. Moreover, the SV19 was polymorphic in both lineages, which could suggest that the SV polymorphism has been present as standing variation in European plaice since at least the time of divergence between the two glacial lineages. This hypothesis was confirmed by the deep haplogroup divergence observed in both SV phylogenies. For the shortest branch, the split of the main haplotypes was estimated to 200 kya (470 kya for the longest branch). Although these split dates should be interpreted with caution since they do not take effects from selection or recombination between haplogroups into account, the differences of an order of magnitude between the age of the populations from the continental shelfs and the age of the alleles and the fact that both alleles for one SV are present in both glacial lineages suggest that the SVs are older than the current populations in which they segregate.

#### Introgression of the structural variants

There is growing evidence that ancient polymorphisms may act as the fuel for adaptive divergence (reviewed in Wellenreuther and Bernatchez, 2018, Marques *et al.*, 2019). In several cases, new alleles originate from adaptive introgression from sister taxa, such as observed in the Heliconius butterflies (The Heliconius Genome Consortium *et al.*, 2012), bringing a “supergene” already adapted to specific environmental conditions into a new genetic background (Jay *et al.*, 2018). In order to test this hypothesis of introgression as a possible source of the observed SVs, we used the European flounder, a euryhaline species adapted to low salinity and known to hybridize with the European plaice, as a candidate for the source of the SVs. Under this hypothesis, the introgressed haplogroup in plaice should be less divergent from flounder than the genome on average. However, we found the opposite pattern, with each haplogroup being more than or as divergent from the European flounder as the plaice-flounder divergence inferred from outside the SVs. Thus, a potential flounder origin of SVs was rejected by the phylogeny. Introgression was also rejected by the f4 statistics showing limited evidence of mismatches between the phylogenetic tree and individuals SNP topologies. More data is necessary to assess if the few mismatches that were observed are due to a random process of allele sorting or represent a case of more ancient introgression, pre-dating the formation of the SVs. Hence, our data suggest that the SVs originated after the split of European flounder and European plaice (9Mya). However, other potential introgression sources, more closely related to plaice than flounder, cannot be ruled out. For instance, introgressive hybridisation could have involved “ghost” species/populations which are now extinct or another species that we did not sample in our study. Further analyses should thus be performed to fully understand the origin of the reported SVs in European plaice.

### The population structure in European plaice and the contribution of the SVs

#### Population structure in European plaice

As mentioned above, we have confirmed the isolation of the Icelandic population which has also been described in previous work (Hoarau *et al.* 2002). It has been hypothesized that this differentiation and the genetic diversity data of the Icelandic population (lower number of microsatellite alleles) could reflect effects from the isolation on the edge of the distribution area and/or a recent bottleneck in the population (Hoarau et al., 2002). Hoarau *et al.* (2002) suggested that these differences were maintained by the deep oceanic regions separating the continental shelf and Iceland acting as a physical barrier to gene flow. Our analyses provide additional information about the history of the Icelandic population, as the demographic analyses performed here suggest that the Icelandic population represents an old population, established from a different glacial refugium that diverged approximately 112 kya from the other populations of European plaice sampled in this study. This scenario would also be consistent with the fact that observed heterozygosity in the Icelandic population reported with both microsatellite data (Hoarau et al. 2002) and in this study, where diversity was in fact somewhat higher for Iceland than in most of the other populations postglacially established along the distributional edge (Barents Sea and Baltic Sea). In fact, Iceland itself may have been the glacial refuge where a relatively high diversity has been preserved (Maggs *et al.*, 2008). Nevertheless, the physical barrier represented by deep oceanic regions may still be an important factor for maintaining the genetic differences that have evolved during the last glacial maximum.

The reductions of diversity found from the North Sea to the Baltic Sea and from the North Sea to the Barents Sea were also reported by the previous studies with 6 microsatellite loci (Hoarau *et al.*, 2002; Was *et al.*, 2010). However, the numbers of markers used at the time were not sufficient to reliably detect any population differences within this geographical region. The populations studied here were sampled along the continental shelf coast lines, resembling a stepping-stone model of isolation, which represents an ideal condition to lead to patterns of isolation-by-distance (Kimura and Weiss, 1964). However, other processes may also be involved in maintaining this pattern (Jenkins *et al.*, 2018), such as the effect of living on the edge of the species distribution range after a post-glacial recolonization. In this case, the general assumption is that colonization of a new habitat is initiated by only a subset of individual from a source population, which increases the effects of genetic drift, leading to the loss of genetic diversity and increase of genetic differences between populations (Hewitt, 2000). As both Barents Sea and Baltic Sea populations are found on the edge of the plaice distribution range and result from two independent events of post-glacial colonization (Hoarau *et al.*, 2002), the genetic differences established during the post glacial recolonization could be maintained by limited gene flow from the population localized in the centre of the distribution (Johannesson and André, 2006), which may be important for maintaining the clear pattern of isolation-by-distance described here. Similar patterns have been reported for various species in association with the salinity gradient of the Baltic Sea (Johannesson and André, 2006, Cuveliers *et al.*, 2012), and along the South-North coast of Norway in taxa with lower dispersal capacities than European plaice (Hoarau *et al.*, 2007; Morvezen *et al.*, 2016).

#### Implication of structural variants for population structure of European plaice

In contrast to genome-wide patterns of weak population structure, the SVs were responsible for population differences much higher (mean *F*_ST_ on haplogroup frequencies over all pairwise comparisons for SV19 = 0.23 and SV21 = 0.12) than the average genome-wide *F*_ST_ (estimated at 0.01). Haplogroup 1, identified as the derived alleles of the SVs (with the lowest diversity and the highest divergence), reached near-fixation along the environmental gradient of the Baltic Sea in both SVs (*f*_SV19 derived_ = 0.94 and *f*_SV21 derived_ = 1.0). However, the same allele was increasing in frequency towards the northern edge of the plaice distribution, in the Barents Sea (*f*_SV19 derived_ =0.52 and *f*_SV21 derived_ =0.97), which was the most distant site from the Baltic Sea. This variation in allele frequency resulted in a geographical structure of the SVs which was different from the genome-wide population structure. The distributional edge of the plaice also represents a common feature associated with the increase of the derived SV allele frequencies (Figure 3a-b), potentially resulting in allele surfing effects in the marginal populations (Excoffier and Ray, 2008). However, selection may also explain the increased frequencies of the derived SV alleles towards the distribution edge. Selection acting on the derived allele of the SVs was also supported by substantial net divergence of haplogroup 1, which was approximately 2.4 times higher than the ancestral divergence in both SV phylogenies (ratio between the long branch and the short branch of the phylogeny). This high divergence may also be linked to the accelerated rate of accumulation of deleterious mutations and background selection (Charlesworth, 1994; Cruickshank and Hahn, 2014; Duranton *et al.*, 2018; Perrier and Charmantier, 2018; Faria *et al.* 2019). However, the accumulation of deleterious mutations alone would make it difficult for the derived allele to fix in several population by drift only, especially in species with large effective size (Ohta, 1973), like plaice.

#### Evidence for local adaptation

The derived allele of the SVs occurred under various habitat conditions, ranging from brackish to marine environments and along temperature and daylight/seasonal gradients within the Atlantic. It is possible that these associations are explained by selection along multiple gradients and on several genes within the SVs. In principle, selection on any of the hundreds of genes within the SVs would result in frequency clines along the environmental gradients (Jay *et al.*, 2018). The SNPs carried by the SVs were previously detected as strong candidates for selection and with significant association with the salinity gradient of the North Sea-Baltic Sea transition zone (Le Moan *et al.*, 2019). In the present study, the highest allele frequency differences at the SVs where found along this transition zone. For instance, the ancestral allele for SV19 (haplogroup 3) disappeared in less than 300 km (Figure 3a) despite being highly frequent within the North Sea (*f*_SV19_ = 0.61) and Norway (*f*_SV19_ = 0.83). Here, gene flow likely still affects the rest of the genome where differentiation is very low at this geographical scale (*F*_ST_ Kattegat vs. Belt Sea is not significantly different from 0). Since gene flow is acting homogeneously on the entire genome under pure migration-drift equilibrium, each haplogroup should be found in high frequency (Slatkin, 1987), as observed in the other range margin. Consequently, the steep allele clines identified along the Baltic Sea environmental gradient were established after the postglacial recolonization and may be maintained by (or coupled with) local adaptation (Kirkpatrick and Barton, 2006). Interestingly, the SV19 carried two heat shock protein genes, which have important functions for cellular stress response (Sørensen *et al.*, 2003), and have already been identified as candidate loci for local adaptation of Baltic Sea populations in several other marine fishes (Hemmer-Hansen *et al.*, 2007 in European flounder; Nielsen *et al.*, 2009 in Atlantic cod; Limborg *et al.*, 2012 in Atlantic herring). Moreover, several biological functions seem to be enriched within the structural variants, notably linked to the immune system, which may also be candidates for local adaptation that could be associated with multiple gradients. Further work focusing on the functional consequences of these SVs will help to understand their effects on the biology of the European plaice.

#### Structural variants promoting evolution at two time-frames

‘Evolution at two time-frames’ has been used to coin the process were several ancient alleles that are locally adapted can be quickly reassembled during more recent colonization of similar environmental conditions. This was initially described by van Belleghem *et al.* (2018) to explain the rapid parallel evolution of post-glacial ecotypes of saltmarsh beetle, which have repetitively evolved after the last glacial maximum but for which the divergence of most alleles under divergent selection can be traced back to a singular origin occurring about 190 kya. Our results suggest that similar processes may be involved in shaping the population structure of the European plaice. Indeed, the derived allele frequencies of the SVs were strongly associated with the North Sea – Baltic Sea transition zone that was connected to the Atlantic ocean 8 kya (Le Moan *et al.*, 2019), but the divergence of the alleles was estimated to be at least 25 times older than the age of the Baltic Sea itself. Moreover, the two SVs have similar estimated divergence times and show similar diversity patterns, suggesting that they share a common origin. Although our analyses did not completely clarify this origin, our study highlights an additional mechanism promoting the rapid reassembling of adaptive variation, i.e. the presence of SVs that maintain adapted alleles together by decreasing recombination between adapted and maladapted backgrounds.

## Conclusions and perspectives

The structural variants in European plaice are among a few examples of large structural variants involved in the maintenance of population structure in marine fishes (e.g. Threespine stickleback: Jones *et al.*, 2012; Atlantic cod: Kirubakaran *et al.*, 2016; Capelin, Cayuela *et al.*, 2019; Atlantic salmon: Lehnert *et al.*, 2019; Atlantic herring, Pettersson *et al.*, 2019; Atlantic Silversides, Therkildsen *et al.*, 2019), but may be present in many other species (e.g. Australasian snapper, Catanach *et al.*, 2019; lesser sandeel, Jimenez-Mena et al., 2019; sea horse, Riquet *et al.*, 2019). However, in several of these examples, SVs are associated with genome-wide heterogeneous differentiation in collinear regions of the genome, which can also reflect signatures of complex demographic history and heterogeneous barriers to gene flow due to endogenous incompatibilities in early stages of speciation (Le Moan *et al.*, 2016; Rougemont *et al.*, 2017; Rougeux *et al.*, 2017, van Belleghem *et al.*, 2018). In European plaice, the North Sea − Baltic Sea differentiation is associated with limited differentiation in the collinear regions of the genome (Figure S10) and several complex demographic models provided a poorer fit to the data than an isolation-with-migration model (Le Moan *et al.*, 2019). Hence, the plaice SVs could potentially be an example from a marine organism of the process examined in the theoretical study by Kirkpatrick and Barton (2006), who described the spread of differentially adapted alleles at inversions despite the presence of high gene flow between adapted and maladapted populations. It thus makes the European plaice an interesting species for further studies of the effects of structural variants on colonization, adaptation and population structure of marine species, particularly for the populations living at the edge of the distribution.

Our data show the genomic heterogeneity of the European plaice population structure. While the differentiation outside the structural variants was associated with demographic history and current connectivity patterns, the two SVs showed high variation in allele frequencies and were associated with more complex patterns of structure which may involve effects from selection. The similar divergence and diversity patterns for the two SVs support the hypothesis that the population diversification of the European plaice has been influenced by evolution at two different time-frames (van Belleghem *et al.*, 2018). The first period followed the origin of the two SVs and was estimated to predate the origin of contemporary populations, while the second period was associated with the contemporary distribution of the SVs towards the edge of the plaice distributional area during a postglacial recolonization. Additional experimental work focusing on the potential fitness effect of these structural variants holds exciting perspectives for understanding their evolution and the role they may play in local adaptation and population structuring. Moreover, longer genomic sequencing reads, as provided by PacBio or nanopore technologies, could confirm if these SVs are chromosomal inversions. In addition, deeper sequencing coverage would allow an exploration of evolutionary signatures within the SV alleles that evolved in different geographical contexts. Finally, identifying the functional role of genes within the putative inversions would be a major step towards understanding the implication of these genomic regions for population divergence and local adaptation. Such studies will provide an interesting framework to assess the evolutionary pathways involved in maintaining structure in this species where dispersal should normally limit population divergence.

## Acknowledgments

We gratefully acknowledge Dorte Meldrup for her assistance with the library preparation, and Jarle Mork, Sigbjørn Mehl, Asgeir Aglen, Gróa Pétursdóttir for providing the samples from the coast of Norway, the Barents Sea and Iceland. Additionally, we thank Nicolas Bierne, Pierre-Alexandre Gagnaire and François Bonhomme for their helpful advises in the early stage of this study, Romina Henriques for her comments on the manuscript, and Marjorlaine Rousselle for her advice on phylogenic analyses. Version 5 of this preprint has been peer-reviewed and recommended by Peer Community In Evolutionary Biology (https://doi.org/10.24072/pci.evolbiol.100095). We are grateful to the Peer Community in Evolutionary Biology; Maren Wellenreuther and the three anonymous reviewers for the reviewing and Denis Bourguet for the technical support. This study received financial support from The European Regional Development Fund (Interreg V-A, project “MarGen”) and from the Ørsted Foundation.

## Data availability statement

Individual raw fasta data, analysed vcf file and scripts (including the R scripts used for statistics and figures, and the two custom bash scripts used to extract the fasta file for the phylogeny and to extract the annotation information for the gene enrichment) are available as a Zenodo archive https://zenodo.org/record/3724510. The modified version of δaδi comprising the 3-step parameter optimisation and the code for the tested models are available on https://popgensealab.wordpress.com/dadi-inference/.

## Supplementary material

**Table S1:** Number of RAD-tag and the number of SNPs obtained after each filtration step in the bioinformatic pipeline

| Filtering | Overall |  |  | Southern dataset |  |  | Northern dataset |  |  |
| --- | --- | --- | --- | --- | --- | --- | --- | --- | --- |
|  | TAGs | SNPs | Ind. | TAGs | SNPs | Ind. | TAGs | SNPs | Ind. |
| **stacks output** | 9877 | 54782 | 255 | 20083 | 106107 | 180 | 43728 | 172150 | 75 |
| **missing ind.** | 9877 | 54782 | 234 | 20083 | 106107 | 166 | 43728 | 172150 | 68 |
| **Singletons** | 8588 | 28052 | 234 | 17376 | 57517 | 166 | 35678 | 94098 | 68 |
| **Hwe** | 8587 | 28016 | 234 | 17342 | 56740 | 166 | 35549 | 92706 | 68 |
| **Maf** | 3084 | 4530 | 234 | 7955 | 13923 | 166 | 20912 | 36760 | 68 |
| **Thin** | 3019 | 3019 | 234 | 7429 | 7429 | 166 | 18880 | 18880 | 68 |

**Table S2:** Pairwise genomic differentiation (*F*_ST_) among the European plaice samples with the SVs (above the diagonal) and with SVs removed (below the diagonal) calculated on the overall dataset. The significance levels are represented by * (p-value < 0.05)

|  | Kattegat | Belt Sea | Baltic Sea | North Sea | Norway | Barents Sea | Iceland |
| --- | --- | --- | --- | --- | --- | --- | --- |
| Kattegat | -- | 0.003* | 0.006* | 0.003* | 0.008* | 0.012* | 0.032* |
| Belt Sea | 0.002* | -- | 0.000 | 0.010* | 0.010* | 0.010* | 0.032* |
| Baltic Sea | 0.003* | 0.000 | -- | 0.013* | 0.013* | 0.012* | 0.033* |
| North Sea | 0.002* | 0.005* | 0.005* | -- | 0.009* | 0.014* | 0.033* |
| Norway | 0.006* | 0.007* | 0.007* | 0.005* | -- | 0.005* | 0.029* |
| Barents Sea | 0.011* | 0.010* | 0.011* | 0.010* | 0.004* | -- | 0.033* |
| Iceland | 0.030* | 0.031* | 0.031* | 0.029* | 0.028* | 0.033* | -- |

**Table S3:** Pairwise genomic differentiation (*F*_ST_) among the European plaice samples based on the chromosomes carrying structural variants: chromosome 19 below the diagonal and chromosome 21 above the diagonal. The significance levels are represented by * (p-value < 0.05)

|  | Kattegat | Belt Sea | Baltic Sea | North Sea | Norway | Barents Sea | Iceland |
| --- | --- | --- | --- | --- | --- | --- | --- |
| Kattegat | -- | 0,023* | 0,039* | 0,014* | 0,054* | 0,036* | 0,020* |
| Belt Sea | 0,009* | -- | 0,000 | 0,073* | 0,018* | 0,009* | 0,002 |
| Baltic Sea | 0,034* | 0,006* | -- | 0,098* | 0,018* | 0,010* | 0,005* |
| North Sea | 0,011* | 0,037* | 0,074* | -- | 0,106* | 0,075* | 0,067* |
| Norway | 0,036* | 0,073* | 0,120* | 0,012* | -- | 0,017* | 0,020* |
| Barents Sea | 0,005* | 0,015* | 0,038* | 0,011* | 0,050* | -- | 0,012* |
| Iceland | 0,047* | 0,064* | 0,092* | 0,045* | 0,000 | 0,050* | -- |

**Table S4:** Pairwise differentiation (*F*_ST_) among the European plaice samples based on the DAPC genotypes for SV19 (below diagonal) and SV21 (above diagonal). The high value of *F*_ST_ (>0.1) are highlighted in bold.

|  | Kattegat | Belt Sea | Baltic Sea | North Sea | Norway | Barents Sea | Iceland |
| --- | --- | --- | --- | --- | --- | --- | --- |
| Kattegat | -- | **0.131** | **0.224** | 0.029 | **0.131** | **0.225** | **0.16** |
| Belt Sea | **0.096** | -- | 0.042 | **0.24** | 0.000 | 0.039 | 0.002 |
| Baltic Sea | **0.324** | **0.098** | -- | **0.331** | 0.043 | 0.000 | 0.027 |
| North Sea | 0.067 | **0.287** | **0.533** | -- | **0.243** | **0.331** | **0.268** |
| Norway | **0.312** | **0.564** | **0.764** | **0.114** | -- | 0.000 | 0.041 |
| Barents Sea | 0.022 | **0.195** | **0.425** | 0.023 | **0.254** | -- | 0.031 |
| Iceland | 0.013 | **0.154** | **0.371** | 0.039 | **0.198** | 0.000 | -- |

**Table S5:** Details of the pairwise inferences performed with δaδi for the 5 best fits for each model. The first column shows the tested model, the following 3 columns detail the statistics used for model selection, in order: the likelihood, the AIC and the delta AIC (relative to the best model). The remaining columns show the estimated parameters: ⊖, the effective population size of Norway and Iceland (Ne_Norw_, Ne_Icel_), the migration rate (m_Norw>Icel_, m_Icel>Norw_) the time of split between the populations (T_S_), the time of secondary contact and time of ancestral migration (T_sc_ / T_am_). The best model is highlighted in bold.

| Models | Likelihood | AIC | $\Delta AIC$ | $\Theta$ | $N_{eNorw}$ | $N_{eIcel}$ | $m_{Norw \rightarrow Icel}$ | $m_{Icel \rightarrow Norw}$ | $T_s$ | $T_{sc} / T_{am}$ |
| --- | --- | --- | --- | --- | --- | --- | --- | --- | --- | --- |
| **SI** | 3357 | 6719 | 3509 | 5230 | 0.02 | 0.01 | -- | -- | 0.0004 | -- |
| SI | 3357 | 6719 | 3509 | 5230 | 0.02 | 0.01 | -- | -- | 0.0004 | -- |
| SI | 3357 | 6719 | 3509 | 5230 | 0.02 | 0.01 | -- | -- | 0.0004 | -- |
| SI | 3357 | 6719 | 3509 | 5230 | 0.02 | 0.01 | -- | -- | 0.0004 | -- |
| SI | 3357 | 6720 | 3510 | 5229 | 0.02 | 0.01 | -- | -- | 0.0004 | -- |
| **IM** | 3199 | 6407 | 3197 | 5434 | 0.19 | 0.18 | 38.65 | 39.92 | 0.0185 | -- |
| IM | 3199 | 6407 | 3197 | 5425 | 0.19 | 0.18 | 38.62 | 39.93 | 0.0181 | -- |
| IM | 3199 | 6408 | 3198 | 5447 | 0.19 | 0.19 | 39.38 | 39.94 | 0.0195 | -- |
| IM | 3199 | 6409 | 3199 | 5431 | 0.19 | 0.18 | 39.91 | 38.84 | 0.0182 | -- |
| IM | 3200 | 6410 | 3200 | 5426 | 0.20 | 0.19 | 39.16 | 39.65 | 0.0191 | -- |
| **AM** | 3055 | 6122 | 2912 | 752 | 0.59 | 10.01 | 13.68 | 1.35 | 0.990 | 0.000 |
| AM | 3320 | 6651 | 3441 | 5245 | 0.18 | 0.10 | 0.00 | 59.94 | 0.006 | 0.000 |
| AM | 3327 | 6666 | 3456 | 5272 | 0.17 | 0.16 | 59.97 | 0.00 | 0.008 | 0.000 |
| AM | 3333 | 6678 | 3468 | 5259 | 0.12 | 0.08 | 59.58 | 0.95 | 0.004 | 0.000 |
| AM | 3337 | 6686 | 3476 | 5239 | 0.09 | 0.06 | 0.01 | 59.55 | 0.003 | 0.000 |
| **SC** | **1599** | **3210** | **0** | **1524** | **1.55** | **0.20** | **5.11** | **48.50** | **0.825** | **0.068** |
| SC | 1599 | 3210 | 0 | 1322 | 1.79 | 0.23 | 4.47 | 41.69 | 0.968 | 0.078 |
| SC | 1599 | 3211 | 1 | 1314 | 1.82 | 0.22 | 4.41 | 41.97 | 0.969 | 0.078 |
| SC | 1600 | 3211 | 1 | 1416 | 1.67 | 0.21 | 4.65 | 43.78 | 0.901 | 0.075 |
| SC | 1600 | 3212 | 2 | 1313 | 1.81 | 0.24 | 4.22 | 40.15 | 0.974 | 0.080 |

**Figure S1:**
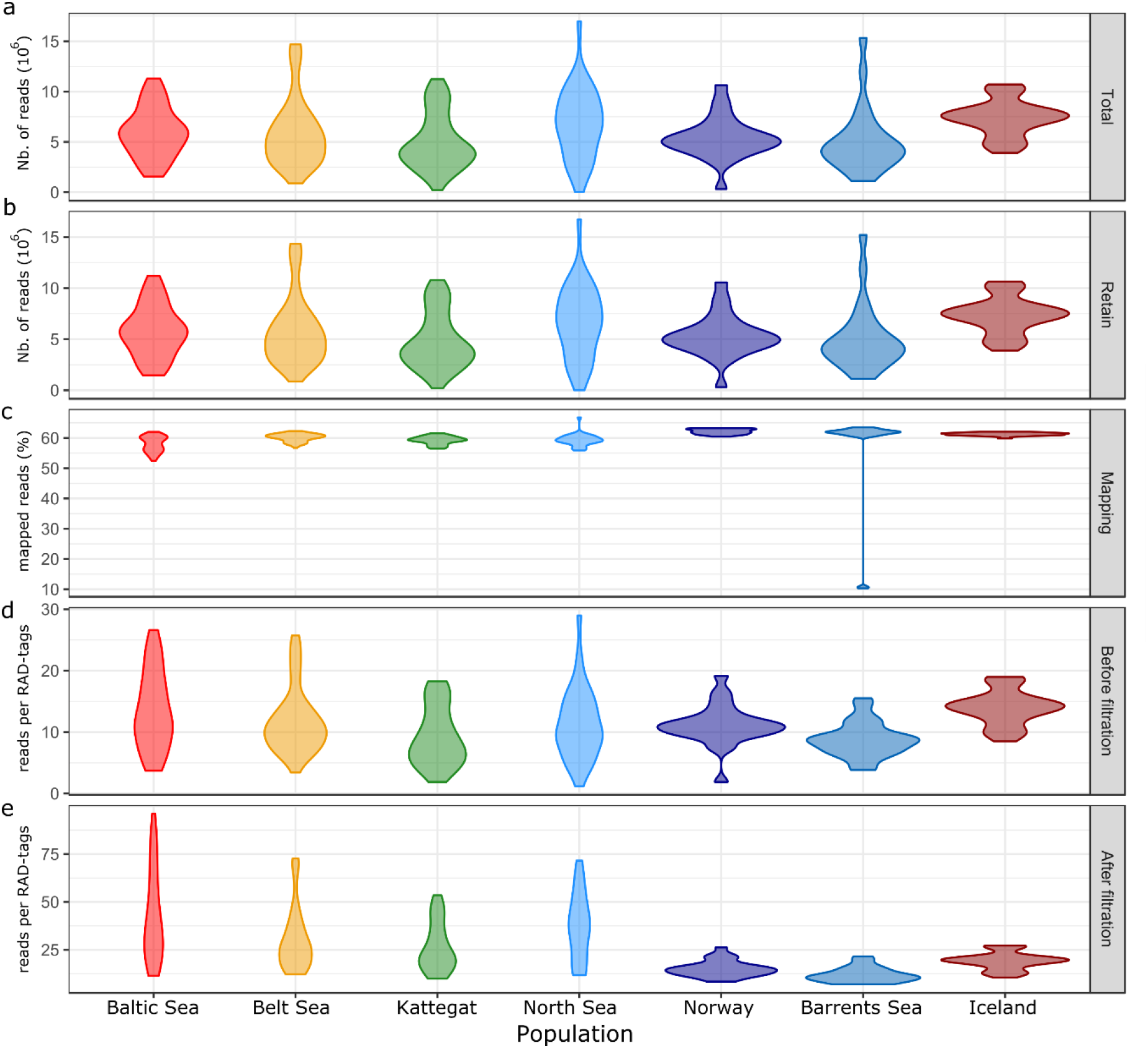
Individual quality statistics of the data per population with a) the total number of reads, b) the number of reads with phred33 quality above 10, c) the percentage of reads mapped back to the Japanese flounder genome, d) the average coverage per RAD-tag and e) the average coverage per RAD-tag after data filtration

**Figure S1:**
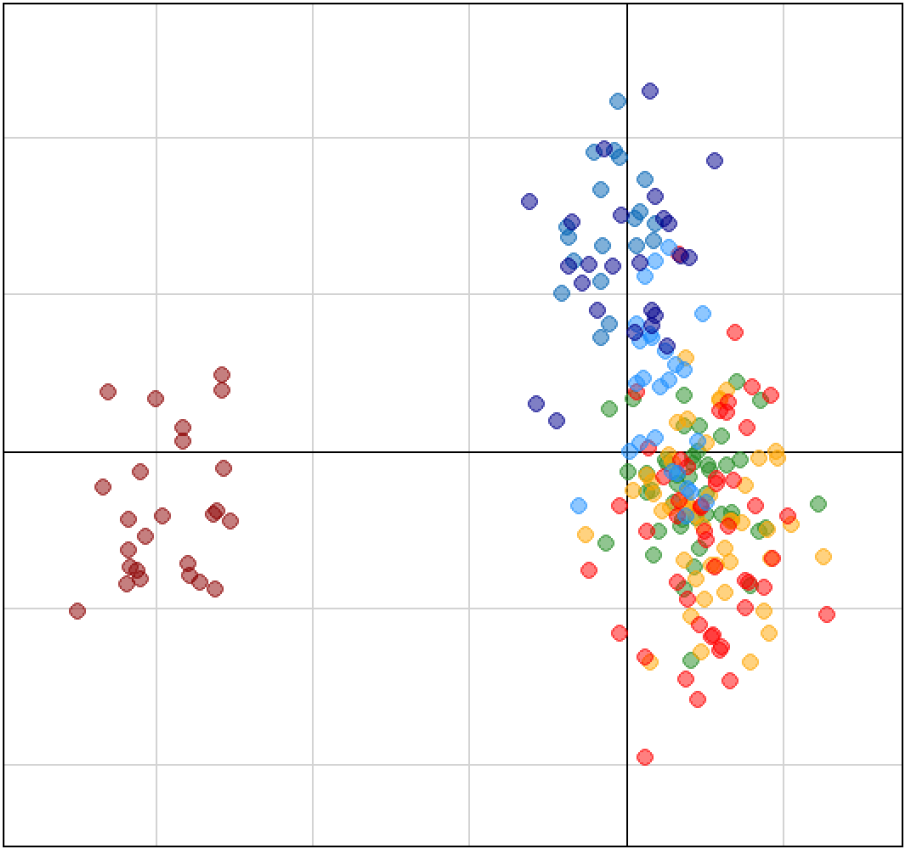
PCA plot based on the 2748 SNPs localised outside the chromosomes carrying SVs. Each dot corresponds to one individuals and the colours correspond to the sampling locations shown on Figure 1a

**Figure S2:**
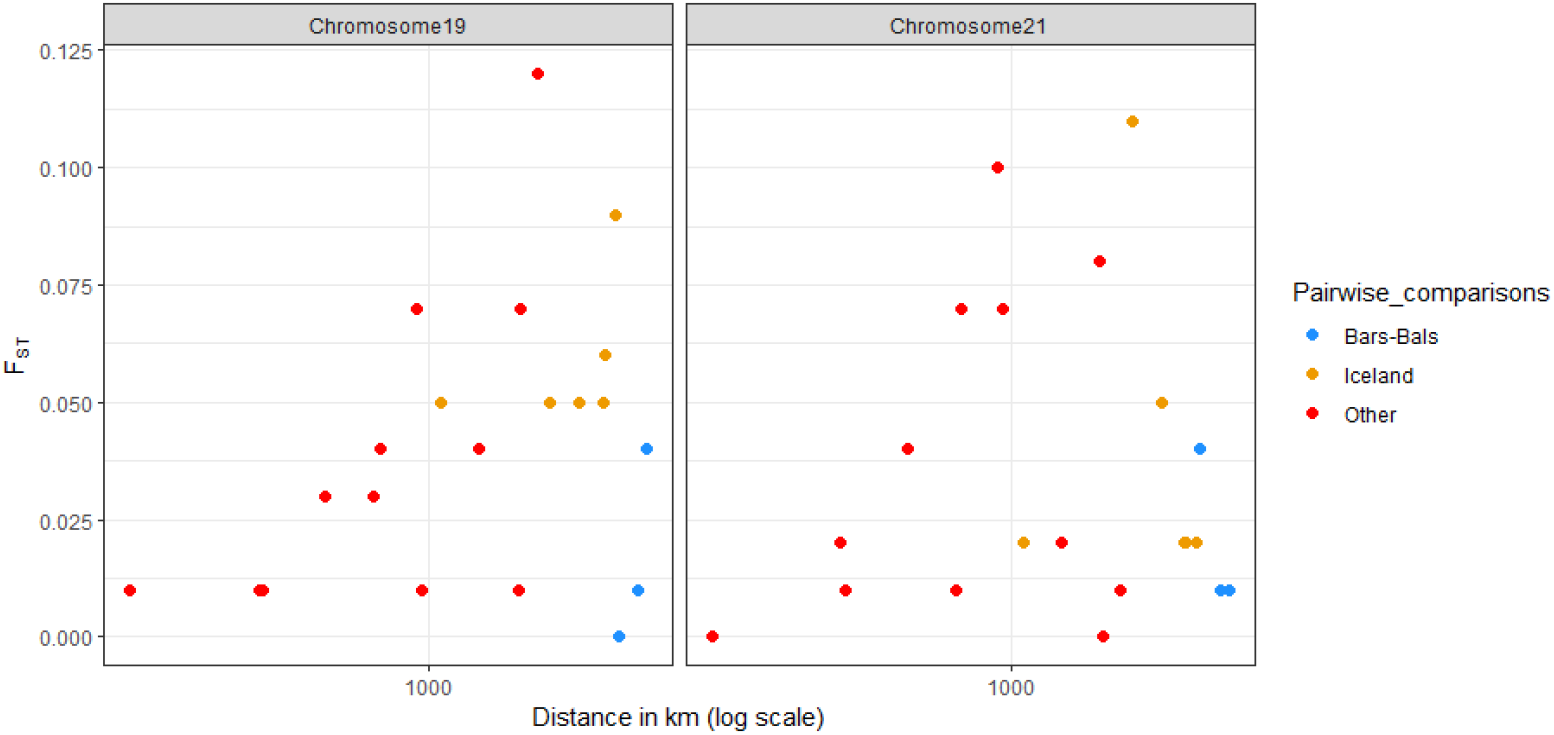
Relationships between *F*_ST_ and geographical distance in European plaice for the polymorphism data of the two chromosomes carrying SVs, chromosome 19 (left) and chromosome 21 (right). Both relations are non-significant (R = 0.06, p = 0.89 for chromosome 19 and R = 0.11, p = 0.35 for chromosome 21)

**Figure S3:**
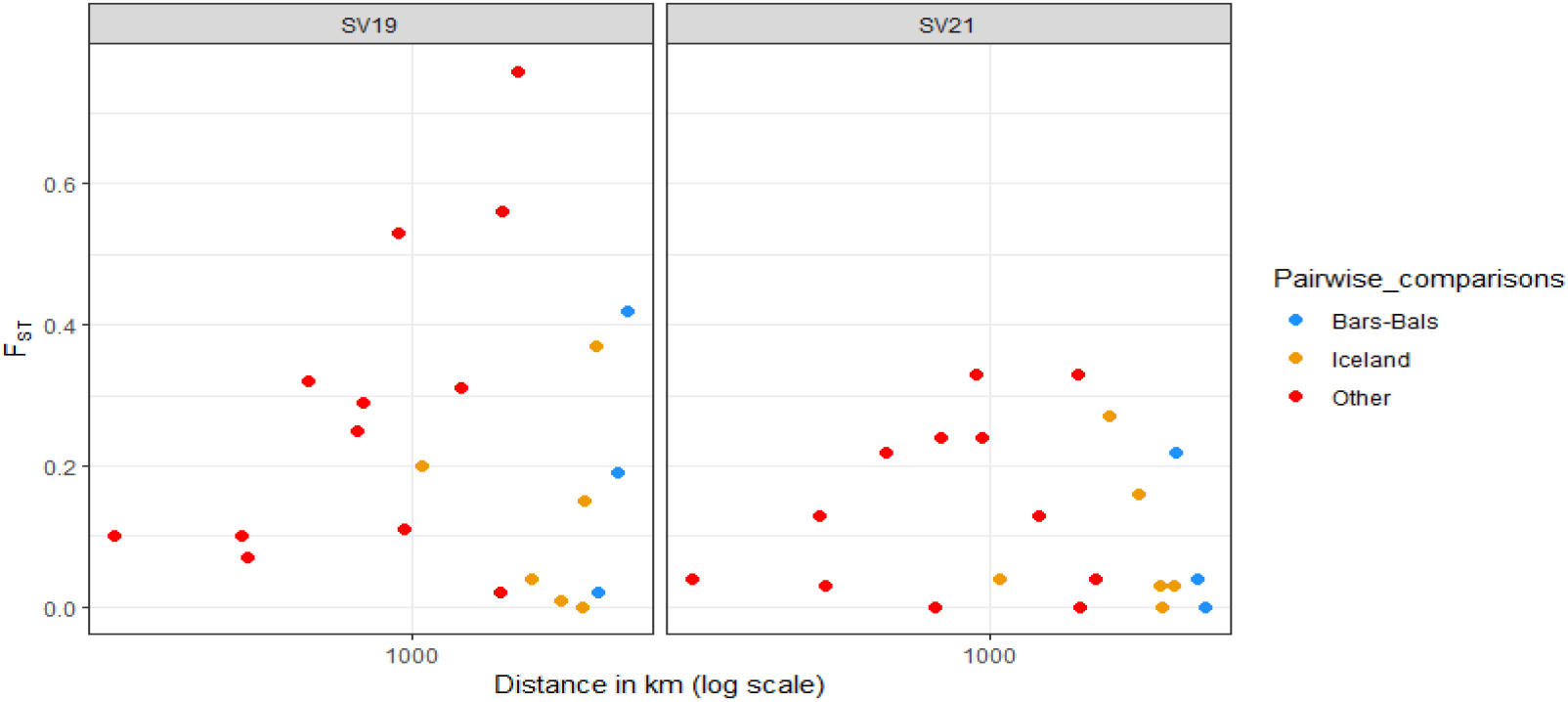
Relationships between F_ST_ for the genotypes of the structural variants SV19 (left) and SV21 (right) inferred from DAPC and geographical distance. Both relations are non-significant (R = 0.00, p = 0.99 for the SV19 and R = −0.23, p = − 0.29, for the SV21).

**Figure S4:**
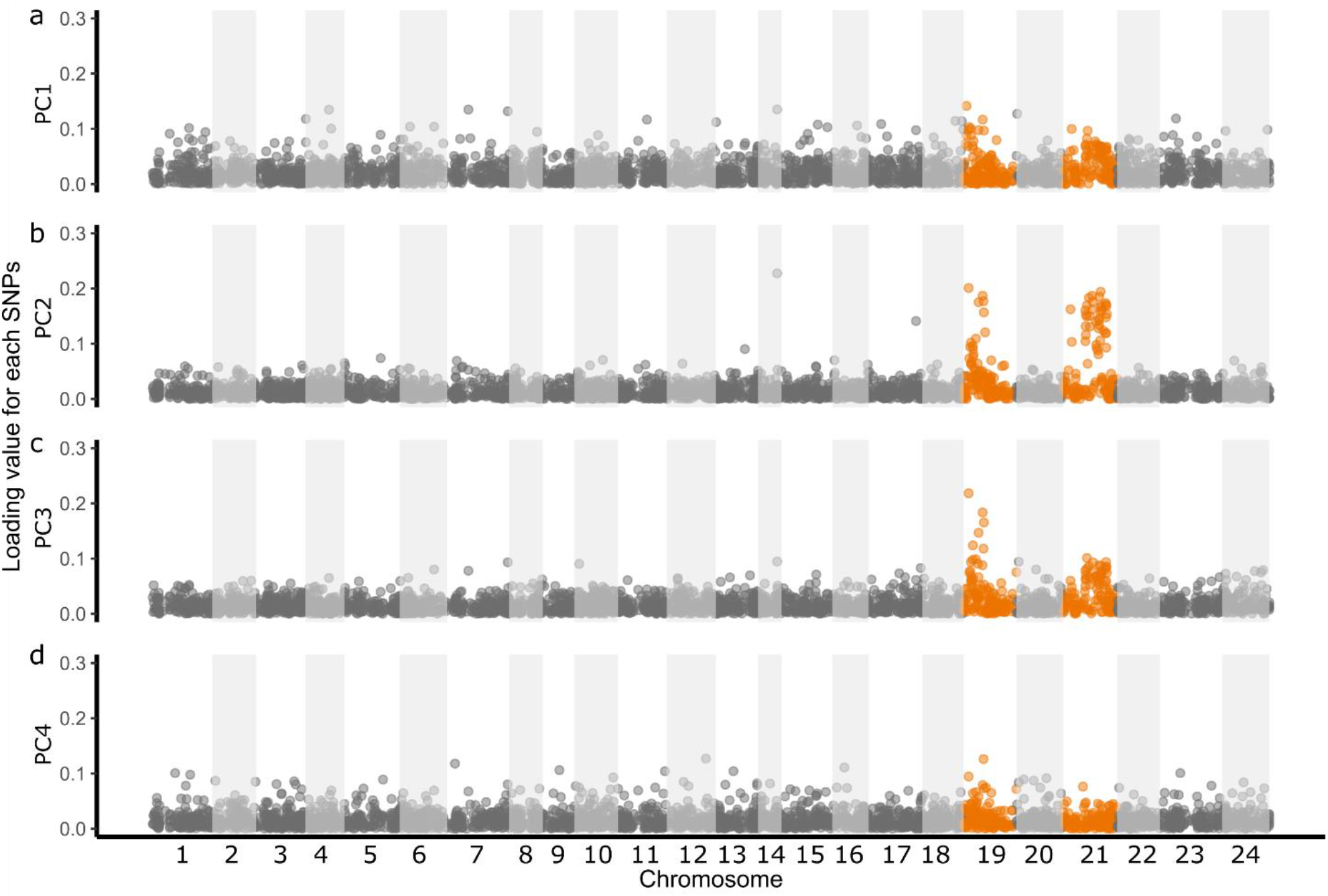
Loading plot of the individual SNP contributions to axes in the PCA analyses. Axis 1 in (a), axis 2 in (b), axis 3 in (c) and axis 4 (d). The orange colour highlights the two chromosomes carrying structural variants.

**Figure S5:**
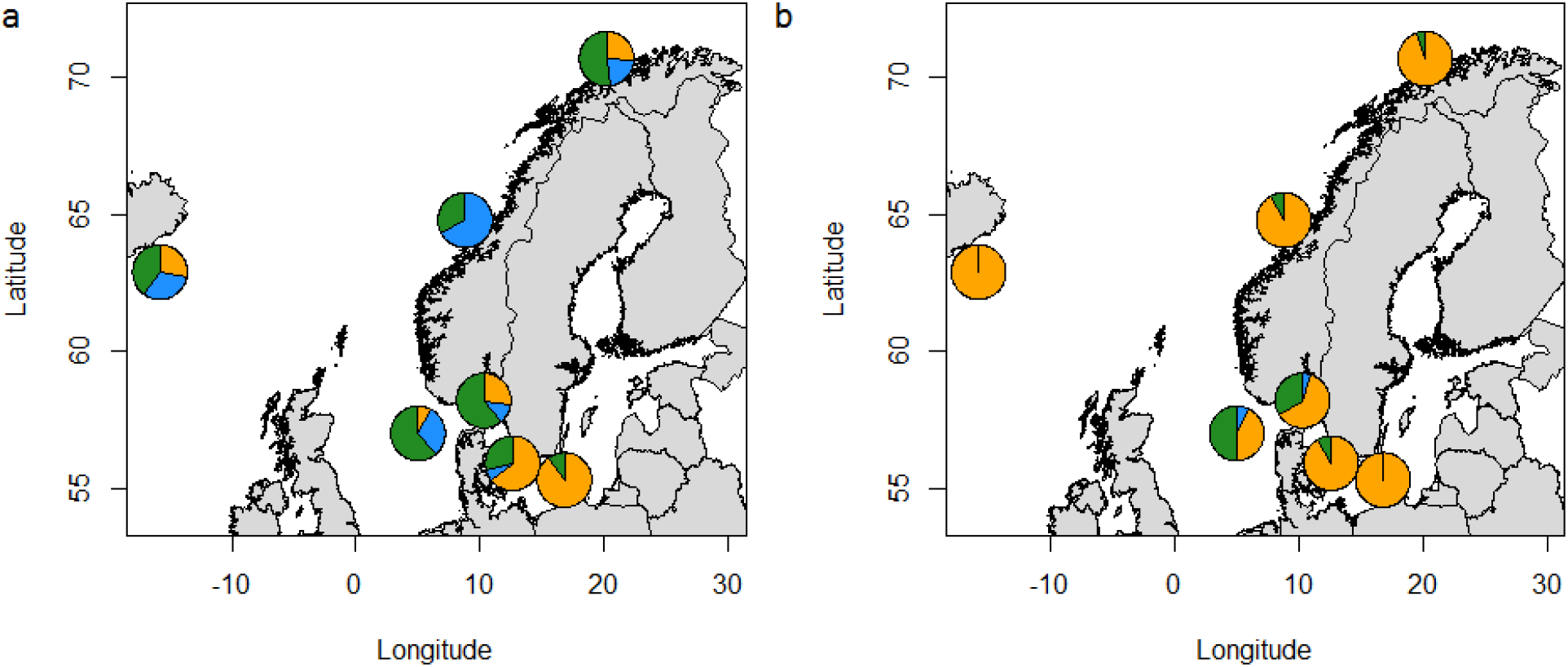
Map of the genotype frequencies across the different sampling sites for SV19 (a) and SV21 (b). Blue represents the frequency of haplogroup 3 (homozygous for the ancestral allele), green represents the frequency of haplogroup 2 heterozygotes and orange represents the frequency of haplogroup1 (homozygous for the derived allele).

**Figure S6:**
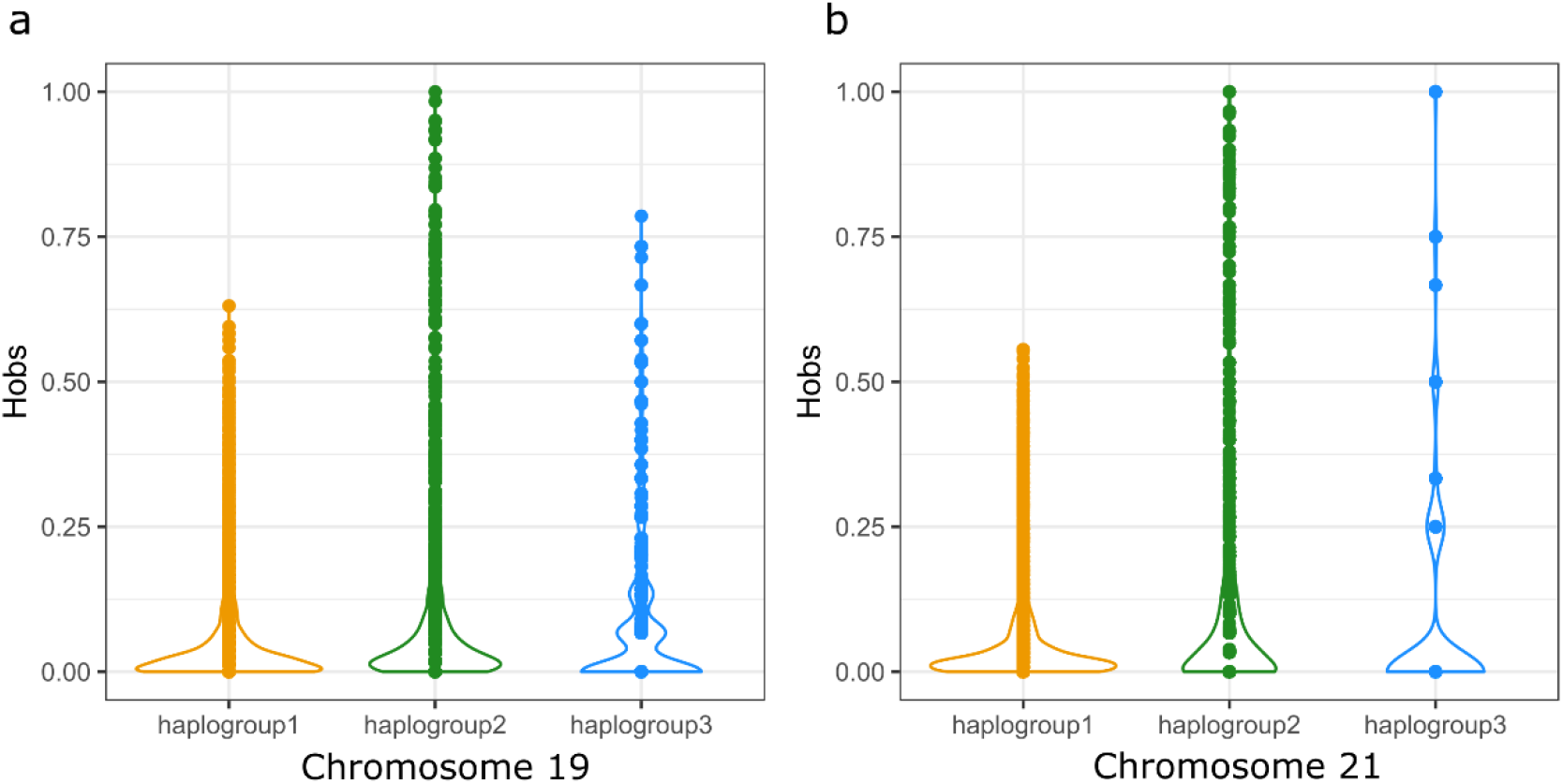
Observed heterozygosity for the DAPC clusters for the chromosomes 19 (a) and 21 (b) carrying SVs. The high variation of the homozygous individuals for haplogroup3 in b is due to the low number of individuals (n=4).

**Figure S7:**
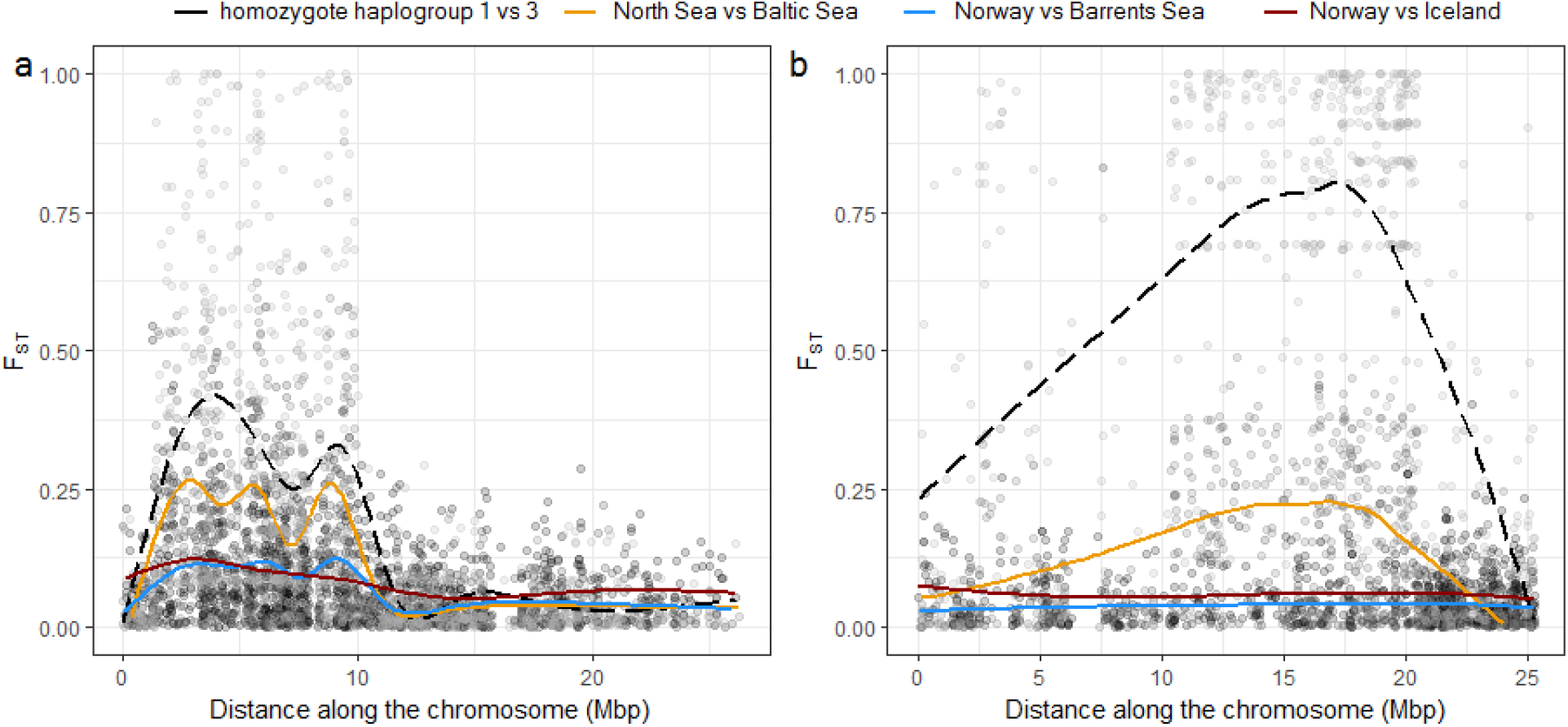
F_ST_ along chromosome 19 in (a) and 21 in (b). Each dot represents the estimate of pairwise F_ST_ for one SNP, and each variation of grey colour corresponds to different pairwise calculations. The line corresponds to the smoothed estimated mean F_ST_ over bins of 10kb (one colour = one comparison)

**Figure S8:**
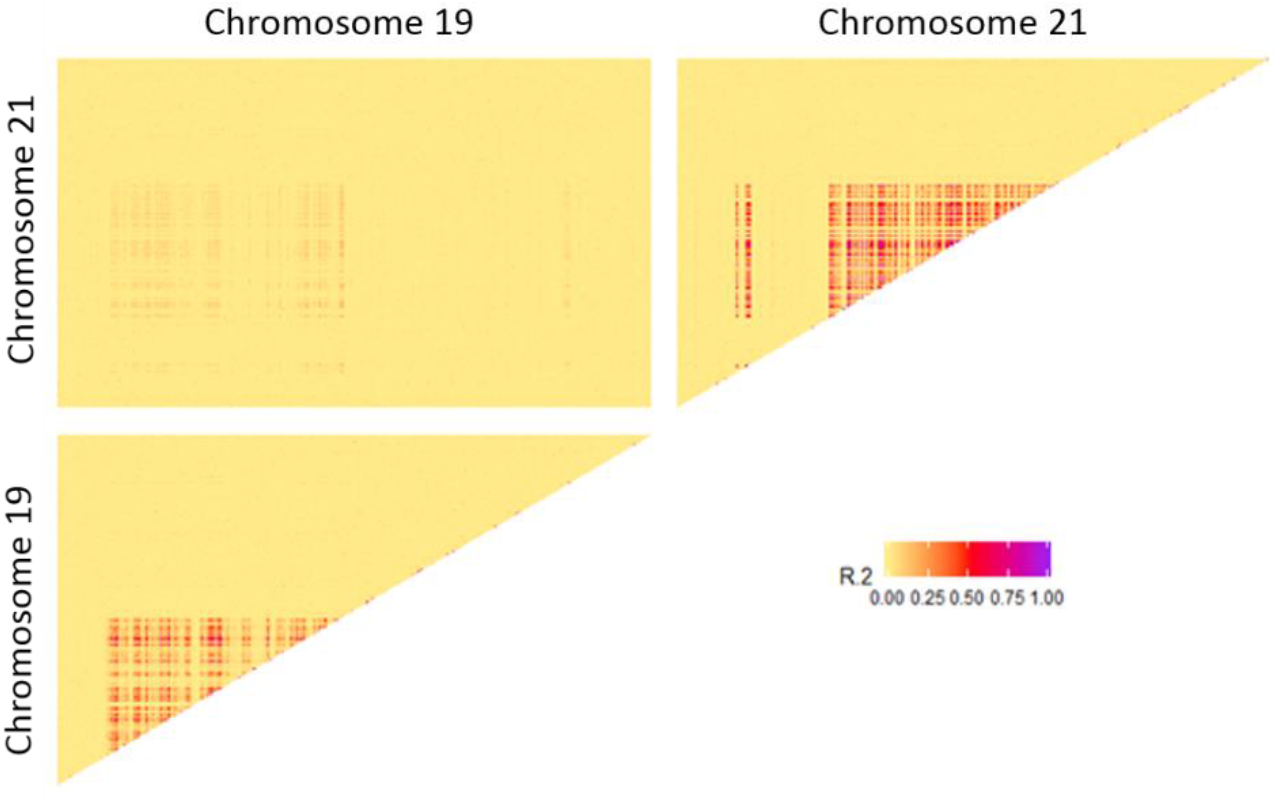
Heatmap of Linkage Disequilibrium between SVs in the European plaice

**Figure S9:**
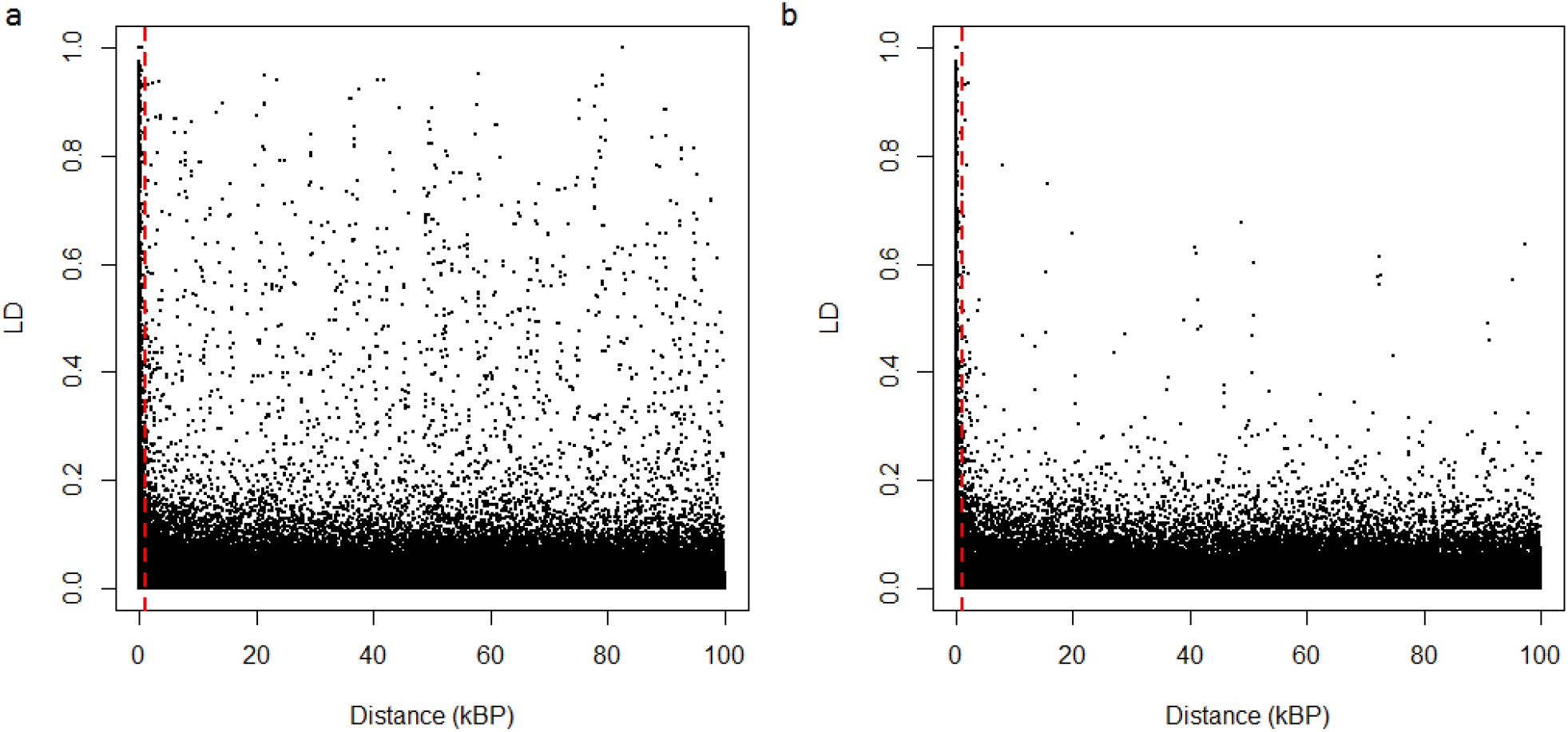
Linkage Disequilibrium decay over 100kb sliding windows when including the SVs (a) and without the SVs (b) of the European plaice. The red line shows the 1kb used as threshold to remove the effect of physical linkage from the demographic and population structure analyses.

**Figure S10:**
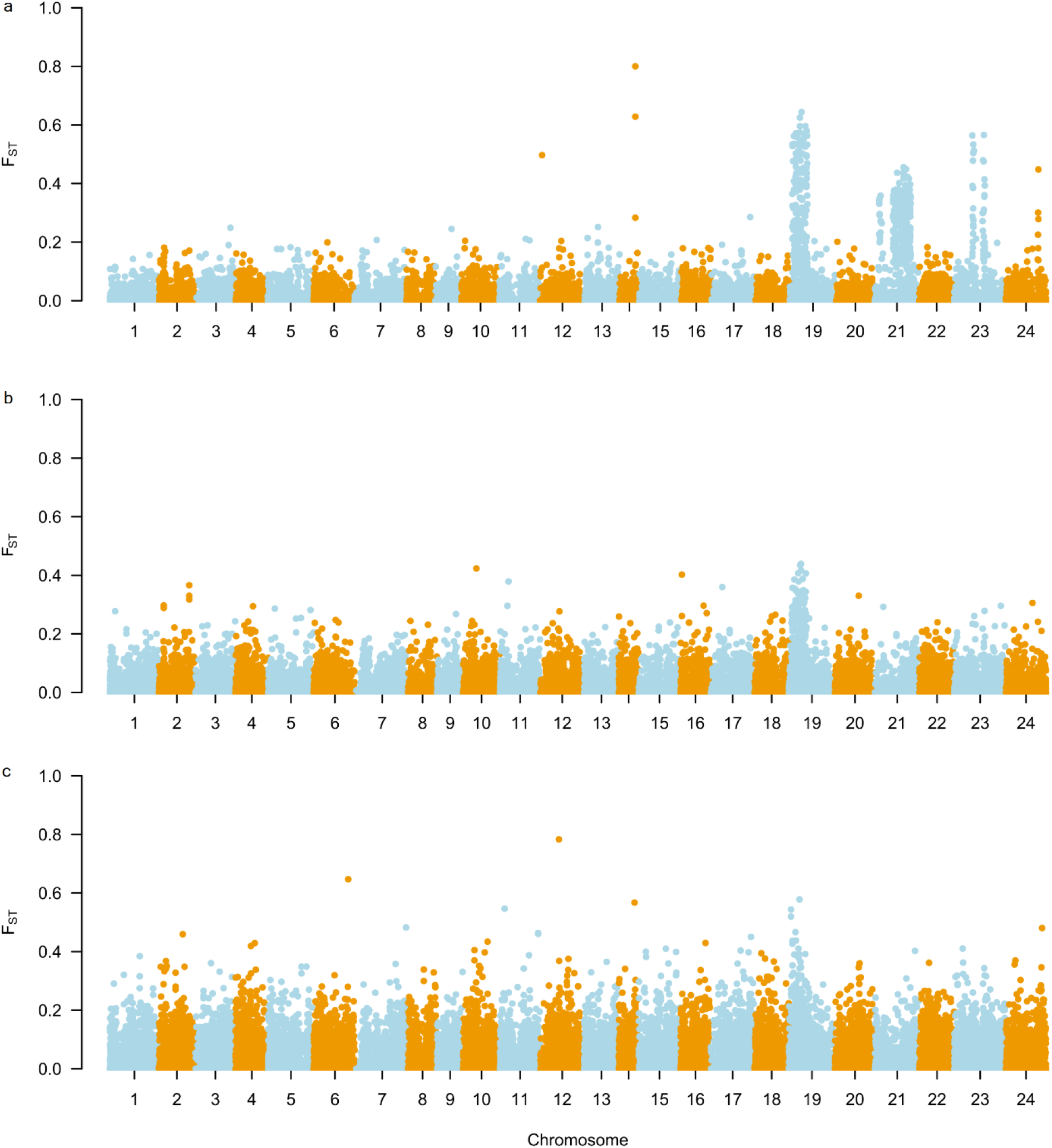
F_ST_ for different pairwise comparisons in the European plaice. North Sea vs Baltic Sea (a), Norway vs Barents Sea (b), and Norway vs Iceland (c). The two structural variants are located on chromosomes 19 and 21.

**Figure S11:**
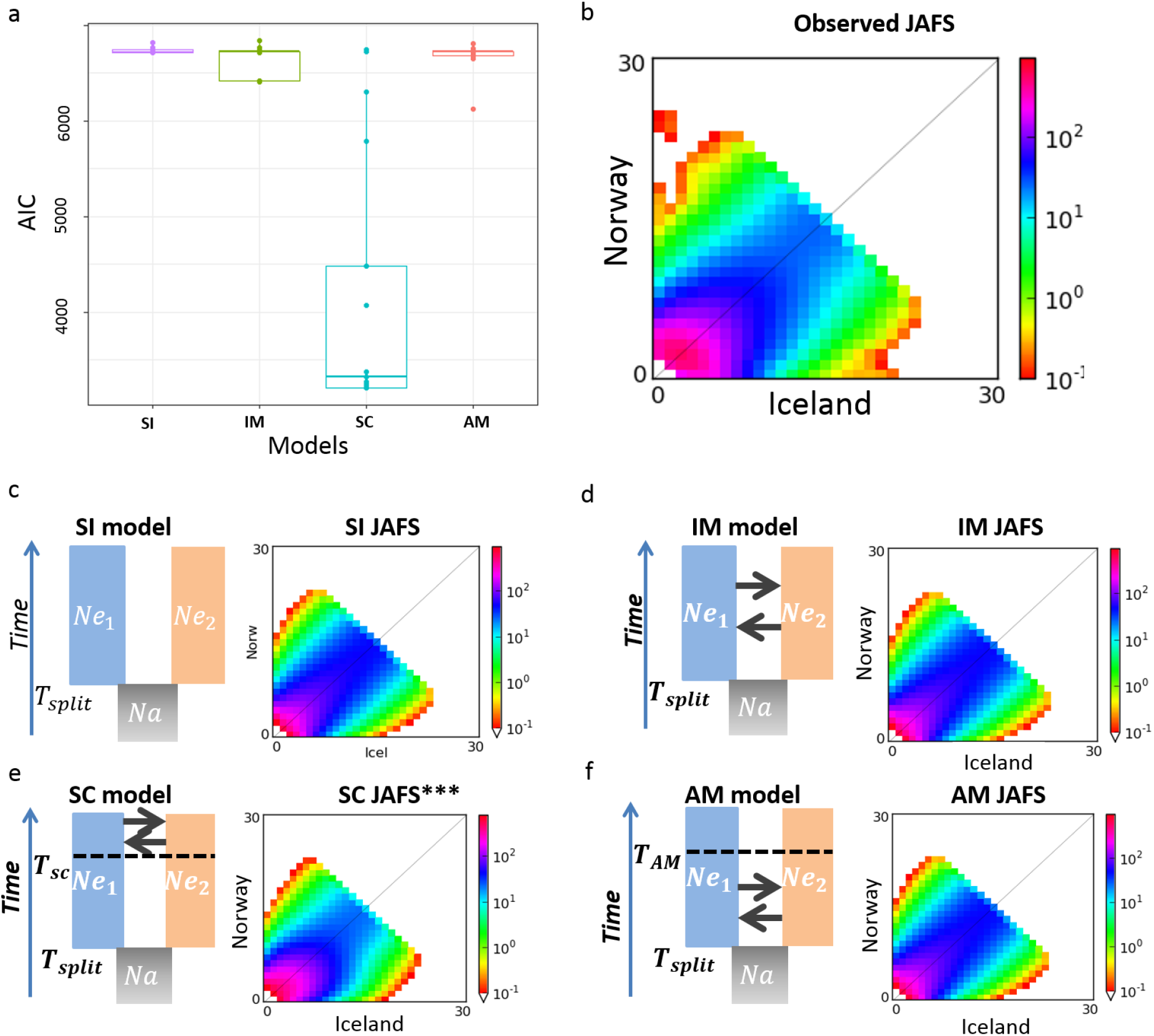
Demographic inferences between Norway and Iceland. a) boxplot of the AIC for the 20 replicates of inferences per model. b) observed joint allelic frequency spectrum (JAFS) between Norway and Iceland. c, d, e and f models SI, IM, SC, AM, respectively, with the theoretical JAFS of the best run for each model.

